# Neuron subtype-specific effector gene expression in the Motor Ganglion of Ciona

**DOI:** 10.1101/641233

**Authors:** Susanne Gibboney, Kwantae Kim, Christopher J. Johnson, Jameson Orvis, Paula Martínez-Feduchi, Elijah K. Lowe, Sarthak Sharma, Alberto Stolfi

## Abstract

The central nervous system of the *Ciona* larva contains only 177 neurons. The precise regulation of neuron subtype-specific morphogenesis and differentiation observed in during the formation of this minimal connectome offers a unique opportunity to dissect gene regulatory networks underlying chordate neurodevelopment. Here we compare the transcriptomes of two very distinct neuron types in the hindbrain/spinal cord homolog of *Ciona*, the Motor Ganglion (MG): the Descending decussating neuron (ddN, proposed homolog of Mauthner Cells in vertebrates) and the MG Interneuron 2 (MGIN2). Both types are invariantly represented by a single bilaterally symmetric left/right pair of cells in every larva. Supernumerary ddNs and MGIN2s were generated in synchronized embryos and isolated by fluorescence-activated cell sorting for transcriptome profiling. Differential gene expression analysis revealed ddN- and MGIN2-specific enrichment of a wide range of genes, including many encoding potential “effectors” of subtype-specific morphological and functional traits. More specifically, we identified the upregulation of centrosome-associated, microtubule-stabilizing/bundling proteins and extracellular matrix proteins and axon guidance cues as part of a single intrinsic regulatory program that might underlie the unique polarization of the ddNs, the only descending MG neurons that cross the midline.

## Introduction

Genetic information is a major determinant of the morphological and physiological properties of individual neurons, as well as the connectivity and function of a nervous system and the properties of its component neurons (Baker et al., 2001; Bargmann, 1993; Manoli et al., 2006). Although these can be influenced by external cues and activity-dependent mechanisms (Thompson et al., 2017; Zhang and Poo, 2001), the clearest evidence for genetic determination of neurodevelopment comes from the stereotyped neural circuits that underlie innate behaviors (Kim and Emmons, 2017; Yamamoto and Koganezawa, 2013), or behavioral phenotypes caused by genetic mutations that result in changes to neuronal cell biology or connectivity (Branicky et al., 2014; White et al., 1992). Although a major focus of modern neuroscience is to dissect behavior at the level of individual genes, neurons, and specific synaptic connections (Luo et al., 2008), we have yet to decipher even the simplest nervous systems. Part of this difficulty stems from the fact that few organisms studied so far have proven tractable enough for the simultaneous investigation of gene function, neuronal activity, circuit connectivity and behavior.

The first synaptic connectivity network, or “connectome” (Sporns et al., 2005) to be fully mapped was that of *Caenorhabditis elegans*, a nematode that has only 302 neurons (White et al., 1986). Also a genetic and developmental model organism, *C. elegans* has delivered many key neurobiology breakthroughs, many of which were prompted by specific insights gleaned from the connectome (Chalfie et al., 1985; Jang et al., 2012). A second connectome, that of the larva of the tunicate *Ciona intestinalis*, was recently completed (Ryan et al., 2016, 2017, 2018). Vertebrates are the sister group to the tunicates within the chordate phylum (Delsuc et al., 2006), and this close genetic relationship has prompted the study of conserved, chordate-specific mechanisms of neurodevelopment in *Ciona* (Nishino, 2018). The central nervous system (CNS) of the *Ciona intestinalis* larva has only 177 neurons (Ryan and Meinertzhagen, 2019), making it the smallest described in any animal. This minimal, but comprehensive, connectome has neatly dovetailed with cell lineage and gene regulatory network studies performed on the *Ciona* species complex (referred to from now on as simply “Ciona”)(Cole and Meinertzhagen, 2004; Horie et al., 2018b; Ikuta and Saiga, 2007; Imai et al., 2009; Nicol and Meinertzhagen, 1988a, b; Sharma et al., 2019). Within this CNS, the development of Motor Ganglion (MG) has been studied in greatest detail. Situated at the base of the tail, just dorsal to the notochord, the neurons of the MG form a simple central pattern that drives the swimming behaviors of the larva (Nishino et al., 2010)(Fig 1A). Of these, a core of only 7 bilaterally symmetric left/right pairs of neurons from the majority of the synaptic connectivity of the MG (Ryan et al., 2016)(Fig 1B), and can all be traced to the A7.8 pair of blastomeres of the 64-cell stage embryo (Cole and Meinertzhagen, 2004; Navarrete and Levine, 2016)(Fig 1C). We refer to these as the “core” MG, as additional cells traditionally assigned to the MG have either been shown to be quite removed from the motor network, e.g. AMG neurons, which serve as peripheral nervous system relay neurons (Ryan et al., 2018), or have yet to be visualized by light microscopy, e.g. Motor Neurons 3 through 5 and MG Interneuron 3 (Ryan et al., 2016).

**Figure 1.**
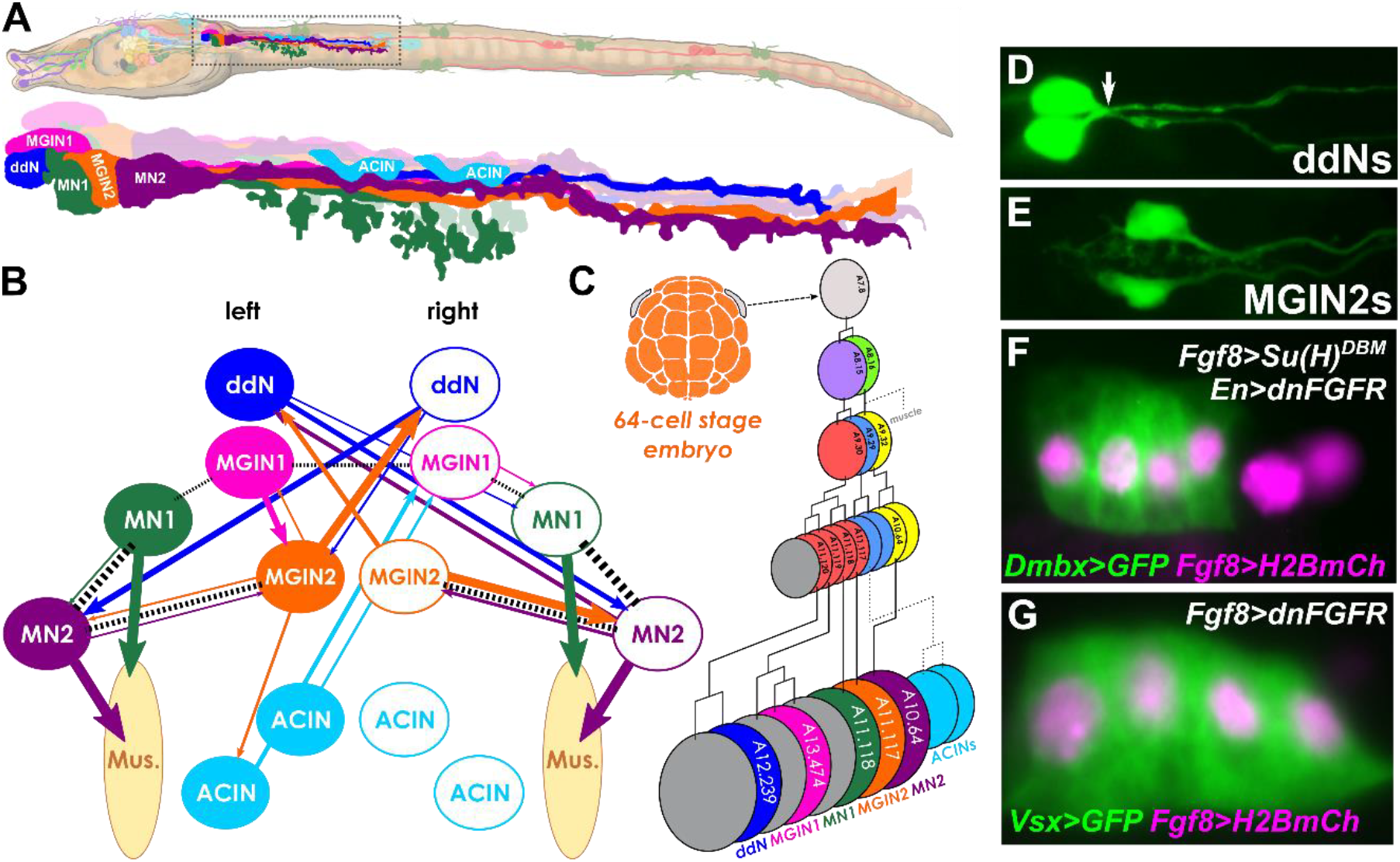
Neuronal subtypes in the Motor Ganglion of Ciona. **A)** Diagram of a *Ciona* larva, with Motor Ganglion (MG) highlighted and magnified in inset. MG neuron morphologies are drawn according to Ryan and Meinertzhagen (2019). Larva illustration by Lindsey Leigh. **B)** Diagram of core MG connectome, adapted from Ryan et al. (2016). Colored arrows indicate major chemical synapses, dashed lines indicate major gap junctions (electrical synapses). Thickness of lines is proportional to cumulative depth of synaptic contact, except for neuromuscular synapses. Only synaptic connections shown are chemical synapses of cumulative depth > 1 μm, and gap junctions of cumulative depth > 6 μm. ACIN left/right symmetry is portrayed even though original connectome was missing a second ACIN on right side. ddN: descending decussating neuron, MGIN1: MG interneuron 1, MGIN2: MG interneuron 2, MN1: motor neuron 1, MN2: motor neuron 2, ACIN: ascending contralateral inhibitory neuron, Mus.: muscles. **C)** Cell lineage diagram of core MG neurons inferred from the literature. Unresolved cell divisions in ACIN lineage indicated by dashed lines. **D)** ddN pair labeled with *Dmbx* reporter construct, arrow indicating axons crossing the midline. **E)** MGIN2 pair labeled with *Pitx* reporter construct. **F)** Supernumerary ddNs generated by early Notch/late FGF inhibition. This condition was used to isolate ddNs by FACS. **G)** Supernumerary MGIN2s generated by early FGF inhibition, condition used to isolate MGIN2s by FACS. Panels D, E, and G adapted from Stolfi and Levine (2011). Panel F adapted from Stolfi et al. (2011).

Within the core MG, each neuron is uniquely delineated by its invariant lineage, molecular profile, morphology, and synaptic connectivity (Cole and Meinertzhagen, 2004; Ryan et al., 2016, 2017; Stolfi and Levine, 2011). Here we focus on the comparison between two very different MG interneuron types: the descending decussating neuron (ddN) and MG Interneuron 2 (MGIN2). As their name implies, ddNs are the only neurons whose axons cross the midline before descending towards the tail (Fig 1D)(Stolfi and Levine, 2011; Takamura et al., 2010). They receive synaptic inputs from peripheral nervous system (PNS) relay neurons and in turn synapse onto other MG neurons, each in particular forming electrical synapses with their respective contralateral Motor Neuron 2 (MN2)(Ryan et al., 2017). The development and synaptic connectivity of the ddNs support homology with the Mauthner cells (M-cells) of the hindbrain of various fish and amphibian species. M-cells initiate the startle reflex during swimming, suggesting they could be mediating similar unilateral tail muscle flexions (“flicks”) in tunicates, in response to mechanosensory stimuli (Ryan et al., 2017). On the other hand, MGIN2s (Fig 1E) are ipsilaterally-projecting descending interneurons whose most salient morphological trait is their extensive dendritic arborization (Stolfi and Levine, 2011). Like the ddNs, they also form conspicuous electrical synapses with MN2s, but receive synaptic inputs mainly from photoreceptor relay neurons and other interneurons of the brain, where the larval light- and gravity-sensing organs are located. Thus, these two MG neurons subtypes might modulate asymmetric swimming behaviors in response to sensory cues processed by distinct thigmotactic (ddNs) and phototactic/geotactic (MGIN2s) pathways (Kourakis et al., 2019; Rudolf et al., 2019; Salas et al., 2018).

By analyzing and comparing the transcriptional profiles of isolated ddNs and MGIN2s, we identified differentially-expressed transcripts enriched in either neuron type and validated the ddN- or MGIN2-specific expression of several genes by mRNA *in situ* hybridization. We hypothesize that many of the genes thus identified are rate-limiting effectors of their unique morphological and physiological characteristics. More specifically, we identify centrosomal microtubule-stabilizing proteins and extracellular matrix components and guidance cues as potential effectors, all expressed by the ddNs themselves, that together might direct the unique mode of polarization and axon outgrowth that we document in the ddNs.

## Materials and methods

### *Ciona robusta* collection and handling

Adult *Ciona robusta* (*intestinalis* Type A) were collected from Pillar Point Marina (Half Moon Bay, CA) or San Diego, CA (M-REP). Dechorionated embryos were obtained and electroporated as previously described (Christiaen et al., 2009a, b). Sequences of plasmids and *in situ* hybridization probe templates not previously published can be found in the **Supplemental Sequences** file. Fluorescent, whole-mount *in situ* hybridization and immunostaining were carried out as previously described (Beh et al., 2007; Ikuta and Saiga, 2007). Images were captured using Nikon, Leica, or Zeiss epifluorescence compound microscopes.

### Whole embryo dissociation for FACS

Embryos were electroporated with the following combinations of plasmids, previously published and described (Stolfi et al., 2011): 60 µg *Fgf8/17/18>Su(H)^DBM^* + 60 µg *Engrailed>dnFGFR* + 61 µg *Dmbx>Unc-76::GFP* + 58 µg *Twist-related.b>RFP* (to generate and isolate ectopic ddNs = “ddN” condition) and 60 µg *Fgf8/17/18>dnFGFR* + 60 µg *En>lacZ* + 61 µg *Vsx>Unc-76::GFP* + 58 µg *Twist-related.b>RFP* (to generate and isolate ectopic MGIN2s = “IN2” condition). Additional unelectroporated embryos were raised in parallel, for sorted, unlabeled cells from whole embryos (“Whole” condition) Embryos were grown to 15.5 hours post-fertilization (hpf) at 16°C and dissociated in trypsin and Ca++/Mg+-free artificial sea water as previously described (Wang et al., 2018).

### Fluorescence-Activated Cell Sorting (FACS) and microarray profiling

FACS was performed on a Coulter EPICS Elite ESP sorter (Coulter Inc.), as previously described (Christiaen et al., 2008), with GFP+ cells selected and RFP+ cells counterselected. “Whole embryo” cells corresponding to cells dissociated from whole unelectroporated embryos were isolated in parallel. RNA extraction was performed using RNAqueous-micro kit (ThermoFisher) as per manufacturer protocol and analyzed by BioAnalyzer (Agilent). Total RNA amounts calculated to correspond to 1364 GFP+ cells was used to prepare each cDNA sample, normalized based on the GFP+ sample that yielded the lowest RNA concentration. Due to even lower RNA concentrations for GFP-“whole embryo” sorted cells, 833 cells were used for replicates 1 and 2, and 2083 for replicate 3, maxing out the volume of RNA solution allowed for cDNA synthesis. See **Supplemental Table 1** for detailed information about each sample used for cDNA synthesis. We used the Ovation Pico WTA System (NuGen) and the Encore Biotin Module (NuGen) to prepare cDNAs and target probes for microrarray, following the manufacturer’s instructions and as previously described (Razy-Krajka et al., 2014). Microarray hybridization, washing, staining, and scanning were performed on a custom Affymetrix GeneChip (ArrayExpress accession A-AFFY-106) according to NuGen protocols and as previously described (Christiaen et al., 2008; Razy-Krajka et al., 2014).

Raw expression values (e.g.. CEL files) for each probe over three biological replicates for each condition (“IN2”, “ddN”, and “Whole”) were used to normalize and compute probe set expression estimates using the robust multi-chips analysis (RMA) algorithm (RMAexpress software) (Bolstad et al., 2003; Smyth, 2004). RMA estimates are available at https://osf.io/n7vr2/ and **Supplemental Table 2**. RMA estimates were averaged across replicates and then converted to Log2 and pair-wise fold-change comparisons (LogFC) calculated and P-values given by 1-tailed type 1 T-test. Probesets were matched to KyotoHoya (KH) gene models (Satou et al., 2008), or prior gene models when KH gene models did not appear to match a probeset sequence. See **Supplemental Table 3** for our new annotation of correspondences between probesets and gene models.

## Results and discussion

We used fluorescence-activated cell sorting (FACS) to isolate specific MG neuron types from synchronized *Ciona robusta* (*intestinalis Type A)* embryos, allowing us to profile their transcriptomes. We took advantage of different genetic manipulations to convert the majority of MG neurons into either supernumerary ddNs or supernumerary MGIN2s, which normally arise from a common progenitor at the neurula stage, the A9.30 pair of blastomeres of the neural plate (Fig 1A)(Stolfi and Levine, 2011). We previously established that irreversibly inhibiting early Notch and late FGF signaling in the anterior cells of the A9.30 lineage converts their progeny into ectopic ddNs, all expressing the ddN marker *Dmbx* (Fig 1F). In contrast, irreversibly inhibiting early FGF signaling converts the entire A9.30 lineage into ectopic MGIN2s, all expressing the MGIN2 marker *Vsx* (Fig 1G). To generate ectopic ddNs or MGIN2s for fluorescence-activated cell sorting (FACS)-mediated cell type isolation, we recapitulated these perturbation conditions by co-electroporating synchronized embryos with specific combinations of plasmids (see Materials and methods for details). Ectopic ddNs were then isolated based on *Dmbx>GFP* expression, while ectopic MGIN2s were isolated by *Vsx>GFP* expression. In both conditions, co-electroporation with *Twist-related.b(KH.C5.554)>RFP* (Abitua et al., 2012) was used to counterselect mesenchyme cells potentially contaminating of our otherwise pure populations of MG neuron types. Embryos were dissociated at 15.5 hours post-fertilization (hpf) at 16°C. Control “whole embryo” cells were dissociated from un-electroporated embryos and subjected to FACS without selection. Total RNA was extracted from sorted cells, followed by cDNA synthesis and transcriptome profiling by microarray, in three independent biological replicates for each condition (see Materials and methods for details).

cDNA libraries were hybridized to Custom-designed Affymetrix GeneChip microarrays (ArrayExpress accession A-AFFY-106)(Christiaen et al., 2008) to quantify transcripts in sorted ddN, MGIN2, and mixed whole-embryo cells, and calculate the enrichment or depletion of ~21,000 individual transcript models in each cell population (**Supplemental Table 2**). Although the custom-made Affymetrix GeneChip microarrays we used were designed prior to the release of the most recent *C. robusta* genome assembly and associated transcript models (KyotoHoya, or KH)(Satou et al., 2008), we re-linked probesets to KH gene models where possible (see Materials and methods for details). Pairwise comparison of ddN and MGIN2 probeset expression values revealed a list of candidate genes that are differentially up-and/or down-regulated in either MG neuron subtype (**Supplemental Table 2**).

We detected 982 probesets that were significantly enriched (LogFC> 0.6, p<0.05) in the ddNs vs. the MGIN2s, and 1245 probesets significantly enriched in MGIN2s vs. ddNs (LogFC< −0.6, p<0.05). This is a slight overestimation of enriched *genes*, since many genes appear to be represented by more than one probeset, due to imprecise gene annotation. By perusing previously published expression patterns of some of the top differentially-expressed genes, we deduced that our FACS-isolated MGIN2 population contained contaminating cells that we failed to counterselect. MGIN2s appeared to be contaminated with epidermis midline cells, based on the presence of epidermal midline markers *Dlx.c* and *Klf1/2/4 (Imai et al., 2004)* in our top 25 MGIN2-enriched genes. This was likely due to the weak expression of *Vsx>GFP* in the dorsal epidermis midline (**Supplemental Figure 1**). This contamination might explain the slightly higher number of genes enriched in the MGIN2 samples.

We next selected a subset of the top differentially expressed genes in either ddNs (Table 1) or MGIN2s (Table 2) to validate by whole-mount fluorescent mRNA *in situ* hybridization (ISH). Some genes were selected on the basis of their statistically significant enrichment in either population, or based on their potential interest to us as candidates for follow-up functional studies.

**Table 1.**
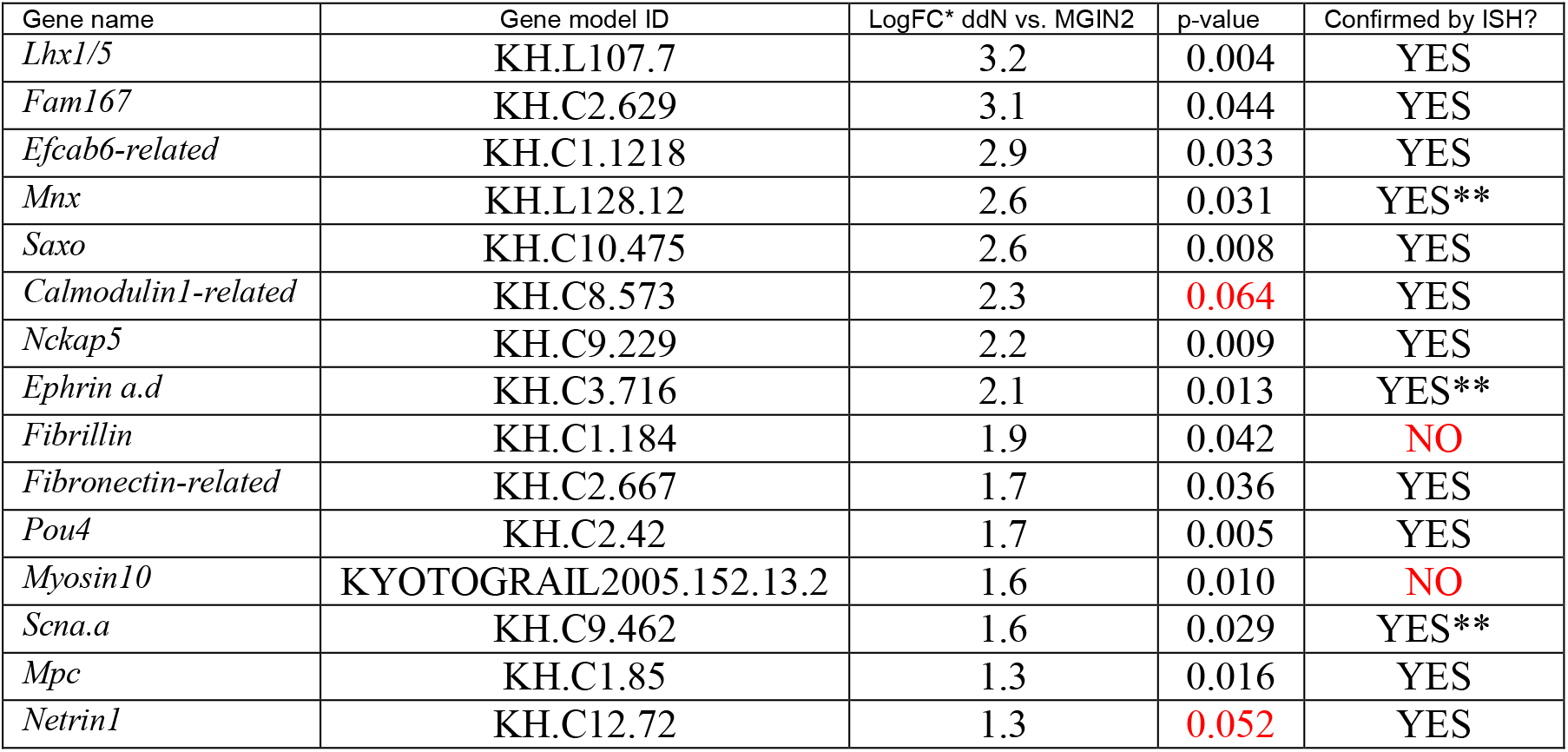
ddN-enriched genes selected for validation by *in situ* hybridization. * for genes represented by more than one probeset, we indicate the highest statistically-significant (p<0.05) LogFC value. Complete dataset available in Supplemental Table 2. ** *Mnx* was enriched in ddN, MN1, and MN2 relative to MGIN2. *Ephrin a.d* and *Scna.a* were enriched in ddN and MN1 relative to MGIN2.

**Table 2.**
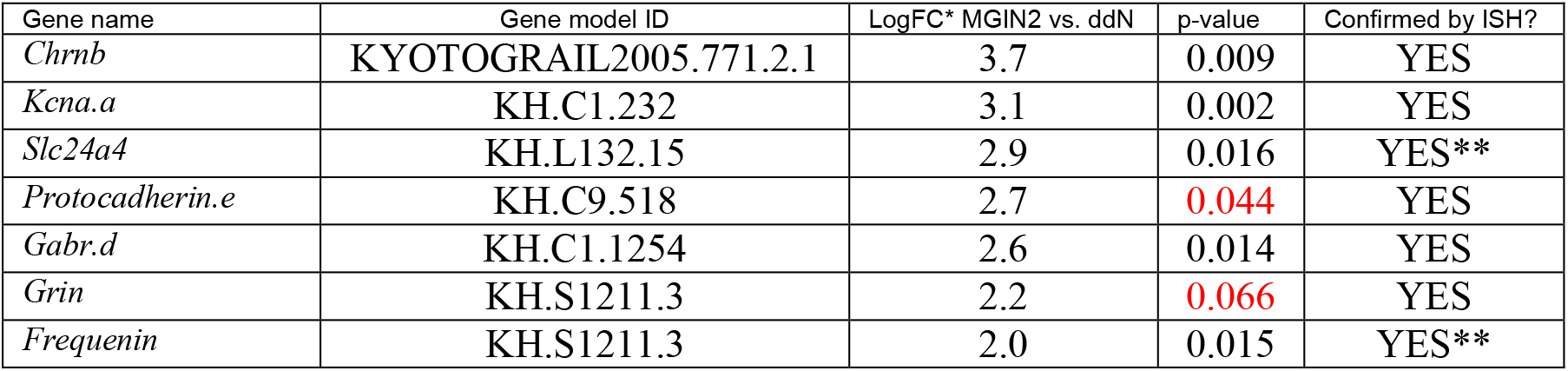
MGIN2-enriched genes selected for validation by *in situ* hybridization. * for genes represented by more than one probeset, we indicate the highest statistically-significant (p<0.05) LogFC value. LogFC values are inverted relative to Table 1 and Supplemental Table 2, to denote enrichment in MGIN2 vs. ddN. ** *Slc24a4* and *Frequenin* were enriched in MGIN2 and MN2 relative to ddN.

| Gene name | Gene model ID | LogFC* MGIN2 vs. ddN | p-value | Confirmed by ISH? |
| --- | --- | --- | --- | --- |
| *ChrnB* | KYOTOGRail2005.771.2.1 | 3.7 | 0.009 | YES |
| *Kcna.a* | KH.C1.232 | 3.1 | 0.002 | YES |
| *Slc24a4* | KH.L132.15 | 2.9 | 0.016 | YES** |
| *Protocadherin.e* | KH.C9.518 | 2.7 | 0.044 | YES |
| *Gabr.d* | KH.C1.1254 | 2.6 | 0.014 | YES |
| *Grin* | KH.S1211.3 | 2.2 | 0.066 | YES |
| *Frequenin* | KH.S1211.3 | 2.0 | 0.015 | YES** |

### ddN-enriched regulators

Our basic strategy was validated by the presence of previously characterized ddN markers among top differentially expressed genes. Although *Dmbx* was indeed the most enriched transcript in the ddNs (LogFC = 6.1), this may have been a result of the probeset detecting portions of the *Dmbx>GFP* reporter that was used to select these cells by FACS. In addition to *Dmbx*, known ddN markers *Lhx1/5* (LogFC = 3.2, Fig 2A)*, Pou4* (LogFC = 1.7, Fig 2B), and *Hox1* (LogFC = 1.6) were among the statistically significant (p < 0.05), top 35 genes most enriched in ddNs relative to MGIN2s. These are all transcription factor-encoding genes and have been previously validated by ISH (Imai et al., 2009; Stolfi et al., 2011). We also found that another transcription factor-encoding gene, *Mnx* (Fig 2C), and an Ephrin signaling molecule-encoding gene, *Ephrin a.d (Efna.d*, KH gene model identifier *KH.C3.716*, Fig 2D) were also highly enriched in the ddNs relative to MGIN2s (LogFC = 2.6 and 2.1, respectively). Expression of *Mnx* and *Efna.d* had not been previously reported in ddNs, but our ISH validation confirmed their expression in the ddN (Fig 2C,D). *Mnx* transcripts were also detected in MN1 and MN2, confirming previous ISH (Imai et al., 2009), while *Efna.d* was also expressed in the sister cell of the ddN (A12.240, which does not give rise to a differentiated neuron in the larval stage), and MN1.

**Figure 2.**
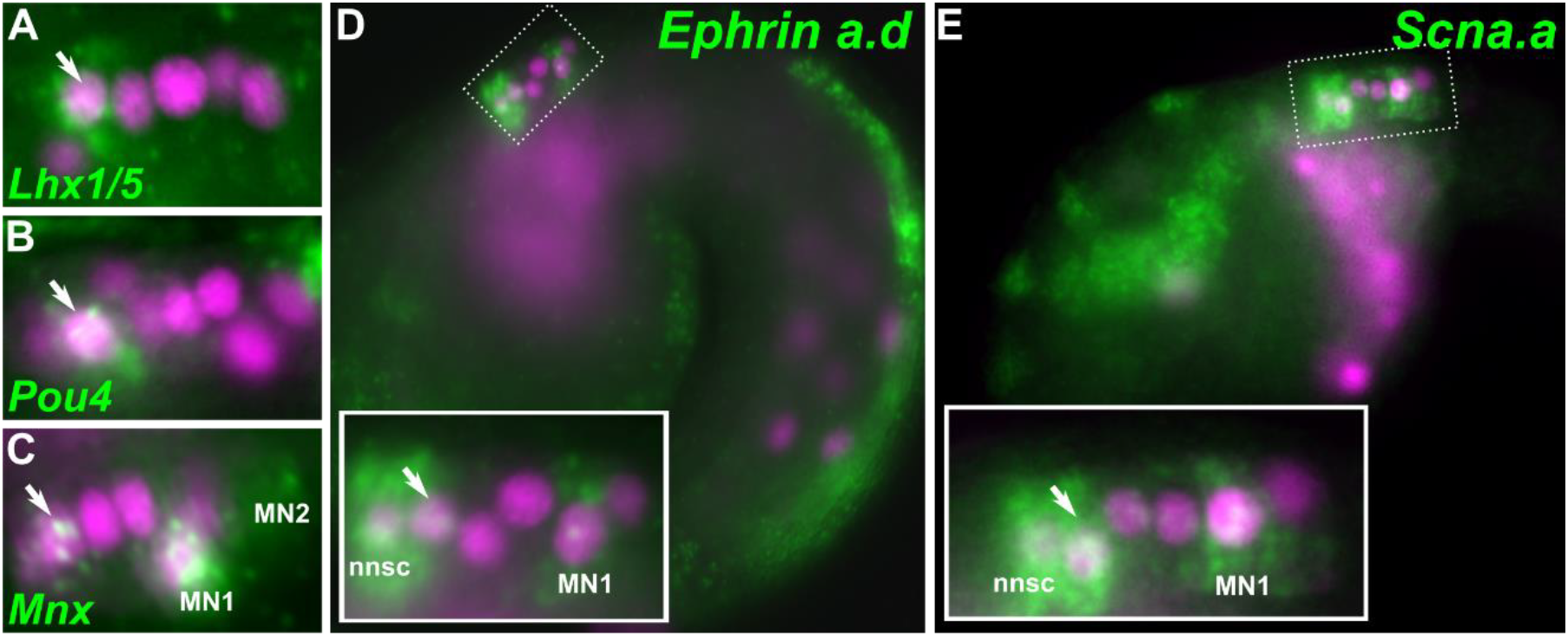
Genes preferentially expressed in ddN vs. MGIN2. Immunostaining of Beta-galactosidase or H2B::mCherry (magenta) in A9.30 lineage progeny nuclei (*Fgf8/17/18* reporters, which also label mesoderm) coupled to *in situ* hybridization (green) of transcripts encoding transcription factors **A)** *Lhx1/5*, **B)** *Pou4*, **C)** *Mnx*, **D)** signaling molecule *Ephrin a.d*, and **E)** voltage-gated sodium channel subunit *Scna.a*. Arrows indicate ddNs. nnsc: non-neuronal sister cell of ddN (A12.240).

Beyond genes encoding transcription factors and signaling molecules, no other potentially ddN-specific transcripts have been previously assayed in detail by ISH. We therefore focused on these, hoping to identify candidate effectors of ddN development and function. For instance, we detected an enrichment for transcripts from the *Sodium voltage-gated channel subunit alpha a* gene *(Scna.a, KH.C9.462*, LogFC = 1.6), encoding a voltage gated sodium channel orthologous to vertebrate NaV1 channels (Katsuyama et al., 2005; Nishino and Okamura, 2018; Okamura et al., 2005). By ISH, we found that *Scna.a* is upregulated specifically in the ddN and in MN1, but not in MGIN2 nor in any other MG neurons at around stage 23 (~15.5 hpf at 16°C), when most are differentiating (Fig 2E). *Ciona* NaV1 (encoded by *Scna.a*) possesses a short, chordate-specific “anchor motif” that in vertebrates is required for dense clustering it the axon initial segment (AIS) through ankyrin-mediated interactions with an actin-spectrin network (Garrido et al., 2003; Hill et al., 2008; Lemaillet et al., 2003). This clustering is crucial for rapid action potential initiation in the proximal axon (Kole et al., 2008). Vertebrate neurons that show dense AIS-specific clustering of NaV1 channels include M-cells and spinal motor neurons (Hill et al., 2008), proposed homologs of ddN and MN1/2, respectively (Ryan et al., 2017). The expression of *Scna.a* in *Ciona* ddN and MN1 suggests that the excitability of these neurons (and ultimately their function) might be similar to those of their vertebrate counterparts, regulated by a conserved, chordate-specific mechanism of subcellular compartmentalization of voltage-gated sodium channels.

### Centrosome-enriched proteins are upregulated in ddNs

Two genes encoding homologs of centrosome-enriched, microtubule-stabilizing proteins were identified among the top ddN-expressed transcripts: *Stabilizer of axonemal microtubules (Saxo, KH.C10.475*, LogFC = 2.6) and *Nck-associated protein 5 (Nckap5, KH.C9.229).* We confirmed the upregulation of both genes in *Ciona* ddNs by ISH (Fig 3A,B). *Saxo* is the sole *C. robusta* ortholog of human *SAXO1* and *SAXO2*, previously known as *FAM154A and FAM154B* respectively. In humans, SAXO1 was found to bind to centrioles and stabilize microtubules (Dacheux et al., 2015). Similarly, *Nckap5* is an ortholog of the closely related human paralogs *NCKAP5* and *NCKAP5L.* In mammals, *NCKAP5L* encodes Cep169, a centrosome-enriched protein that also stabilizes microtubules (Mori et al., 2015a; Mori et al., 2015b). In our profiling, a single probeset detected enrichment of *Saxo*, but fragmented annotation of earlier versions of the *C. robusta* genome resulted in at least 5 independent probesets that we manually annotated as covering the updated *Nckap5* gene model (*KH.C9.229*). These 5 probesets were all significantly enriched in ddNs relative to MGIN2s (LogFC 1.6-2.2, average 1.8).

**Figure 3.**
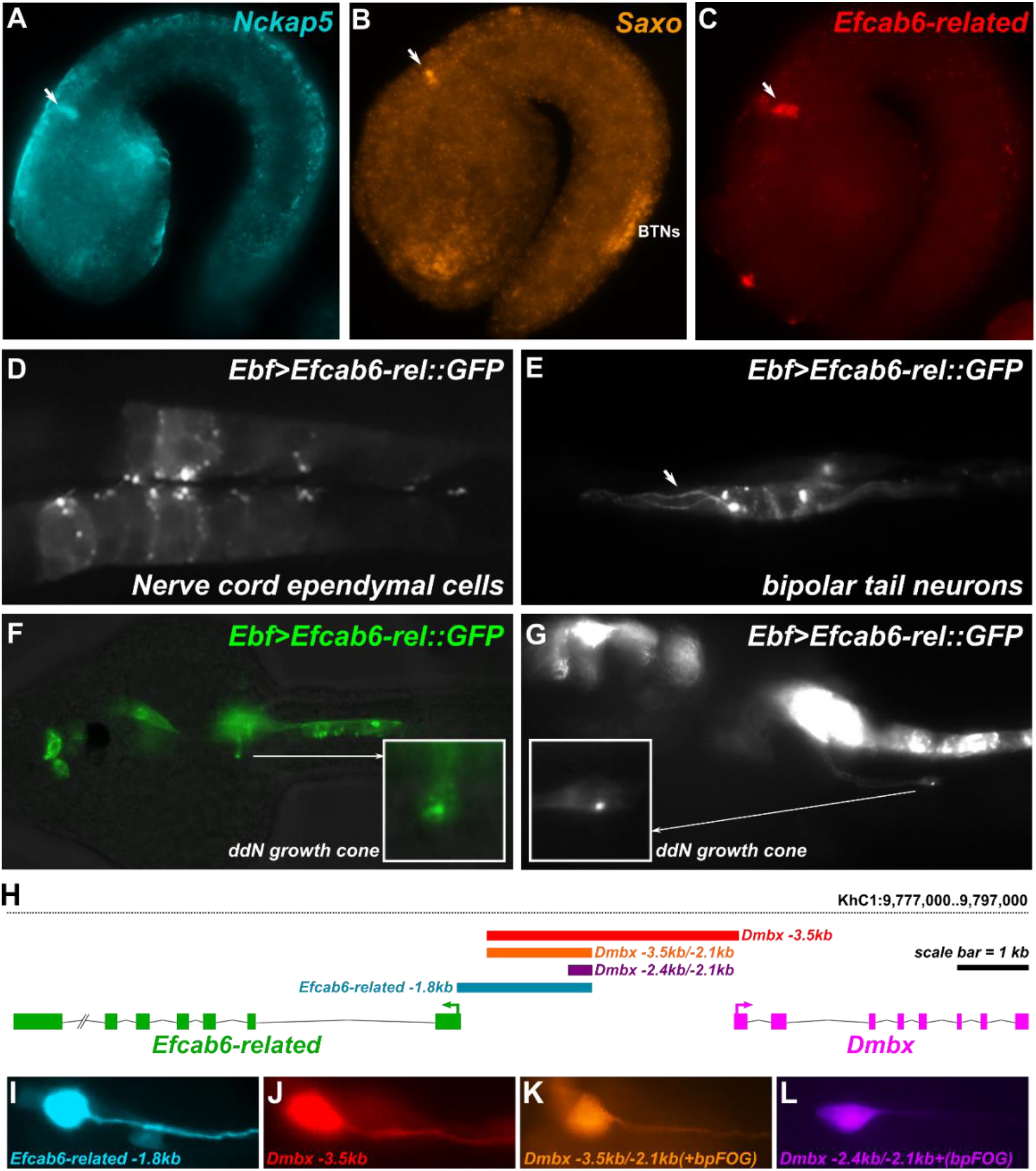
Genes encoding centrosome-localized microtubule-binding proteins are enriched in ddNs. *In situ* hybridization of **A)** *Nckap5*, **B)** *Saxo*, and **C)** *Efcab6-related.* Arrows indicate ddNs. BTNs: Bipolar Tail Neurons. **D)** Efcab6-related::GFP (driven by *Ebf* promoter) labeling presumed basal bodies at apical surface of ciliated ependymal cells of the nerve cord. **E)** Efcab6-related::GFP labeling of centrosomes in differentiating Bipolar Tail Neurons, and labeling microtubule bundles (arrow) in axon. **F)** Efcab6-related::GFP labels a small punctum in the axon growth cone of a ddN as it extends across the midline. **G)** Efcab6-related::GFP punctum also in the growth cone of the ddN axon turning and extending posteriorly down the tail. **H)** Diagram of *Efcab6-related* and *Dmbx* loci and shared *cis*-regulatory sequences. **I-L)** Expression of various *Efcab6-related/Dmbx* reporters in the ddN, color coded according to the diagram in panel H.

In addition to these orthologs of genes encoding previously characterized centrosomal proteins, we also noticed enrichment of *KH.C1.1218* (LogFC = 2.9, Fig 3C), encoding an EF-hand calcium-binding domain-containing protein (See **Supplemental Sequences**). The predicted protein is weakly similar to human EFCAB6 (also known as DJ-1 Binding Protein, or DJBP). However, *KH.L125.4* (and not *KH.C1.1218)* is the predicted ortholog of human *EFCAB6* according to inParanoid (O’Brien et al., 2005). The two *C. robusta* genes appear closely related and might either be descended from a tunicate-specific duplication or from an earlier duplication. *KH.C1.1218* has predicted orthologs in various fish species, *Xenopus tropicalis*, and even platypus, which suggests that this gene may have been lost in certain chordate clades such as placental mammals. For these reasons, we refer to *KH.L125.4* as *Efcab6* and *KH.C1.1218* as *Efcab6-related* from now on.

In human cells, EFCAB6 can inhibit the transcriptional activity of androgen receptor (Niki et al., 2003) through its association with DJ-1, a regulator of oxidative stress response and mitochondrial function (Canet-Avilés et al., 2004; Wang et al., 2012). A mouse knockout line from the Knock Out Mouse Phenotyping Program (KOMP^2^, Jackson Laboratory) for Efcab6 shows inserted reporter gene staining in the developing hindbrain (http://www.mousephenotype.org/data/genes/MGI:1924877), suggesting a potentially conserved role in M-cell/ddN development. Given its seven EF-hand domains, we hypothesized that Efcab6-related might be localized to centrosomes, much like other EF hand-containing, calcium-binding proteins like Centrin or Calmodulin (Ito and Bettencourt-Dias, 2018). Indeed, an Efcab6-related::GFP fusion was specifically localized as puncta near the apical surface of ciliated ependymal cells (Fig 3D), suggesting a centrosomal/basal body localization. In bipolar tail neurons (Stolfi et al., 2015), Efcab6-related::GFP was enriched at the centrosomes and on microtubule bundles in extending axons (Fig 3E). In ddNs, we detected a small punctum of Efcab6-related::GFP in the axon growth cone during extension over the midline (Fig 3F) and later down the tail (Fig 3G). These localization patterns hint at previously unrecognized roles for Efcab6-like proteins in regulating centrosome function, microtubule stabilization, and/or axon extension.

Of further note, *Ciona Efcab6-related* and *Dmbx* are neighboring genes, arrayed in a “head to head” manner and transcribed in opposite directions (Fig 3H). A reporter construct spanning this putative shared *cis-*regulatory module and the translation start site of *Efcab6-related* was sufficient to drive expression in the ddNs (Fig 3I). Since the minimal *cis*-regulatory element that drives Pax3/7-dependent *Dmbx* transcription in the ddNs (Stolfi and Levine, 2011; Stolfi et al., 2011) is roughly equidistant and 5’ to both *Dmbx* and *Efcab6-related* (2.1 kb and 1.9 kb respectively, Fig 3J-L), these two genes likely share a common regulatory element for ddN-specific expression. Given that Dmbx itself is a transcription factor that appears to inhibit proliferation and promote mitotic exit (Stolfi et al., 2011; Wong et al., 2015), this shared *cis*-regulatory element might be essential for coordination of genetically linked, but mechanistically distinct specification and morphogenetic processes in the ddNs.

This ddN-specific expression of these known centrosome-enriched microtubule stabilizing proteins Nckap5 and Saxo, and the previously unrecognized centrosome marker Efcab6-related identified in this study, is interesting given the cellular processes that appear to underlie the unique contralateral axon projection of the ddNs. The ddN axon begins as an initial outgrowth that is oriented towards the neural tube lumen, extending across the midline (Fig 4A). This is immediately preceded by a precisely timed, 180° re-orientation of the intracellular polarity of the cell, as visualized by the position of the Golgi apparatus, starting from an apical position apposing the neural tube lumen to a basal position near the neural tube basal lamina (Fig 4B-D, **Supplemental Table 4**). In all other MG neurons, the Golgi apparatus remain on the apical side (lumen), and the direction of axon outgrowth is instead oriented away from the midline, resulting in an ipsilateral axon trajectory. A similar positioning of the Golgi apparatus on the opposite side of the nucleus relative to the site of axon extension was previously documented in migrating *Ciona* Bipolar Tail Neurons (BTNs), in which a precisely timed, 180° re-orientation of Golgi apparatus position also correlates with the direction of axon extension from an initially anterior orientation to a posterior one (Stolfi et al., 2015). In these cases, the position of the Golgi apparatus is marker for centrosome position, which are tightly linked in chordate cells (Sütterlin and Colanzi, 2010).

**Figure 4.**
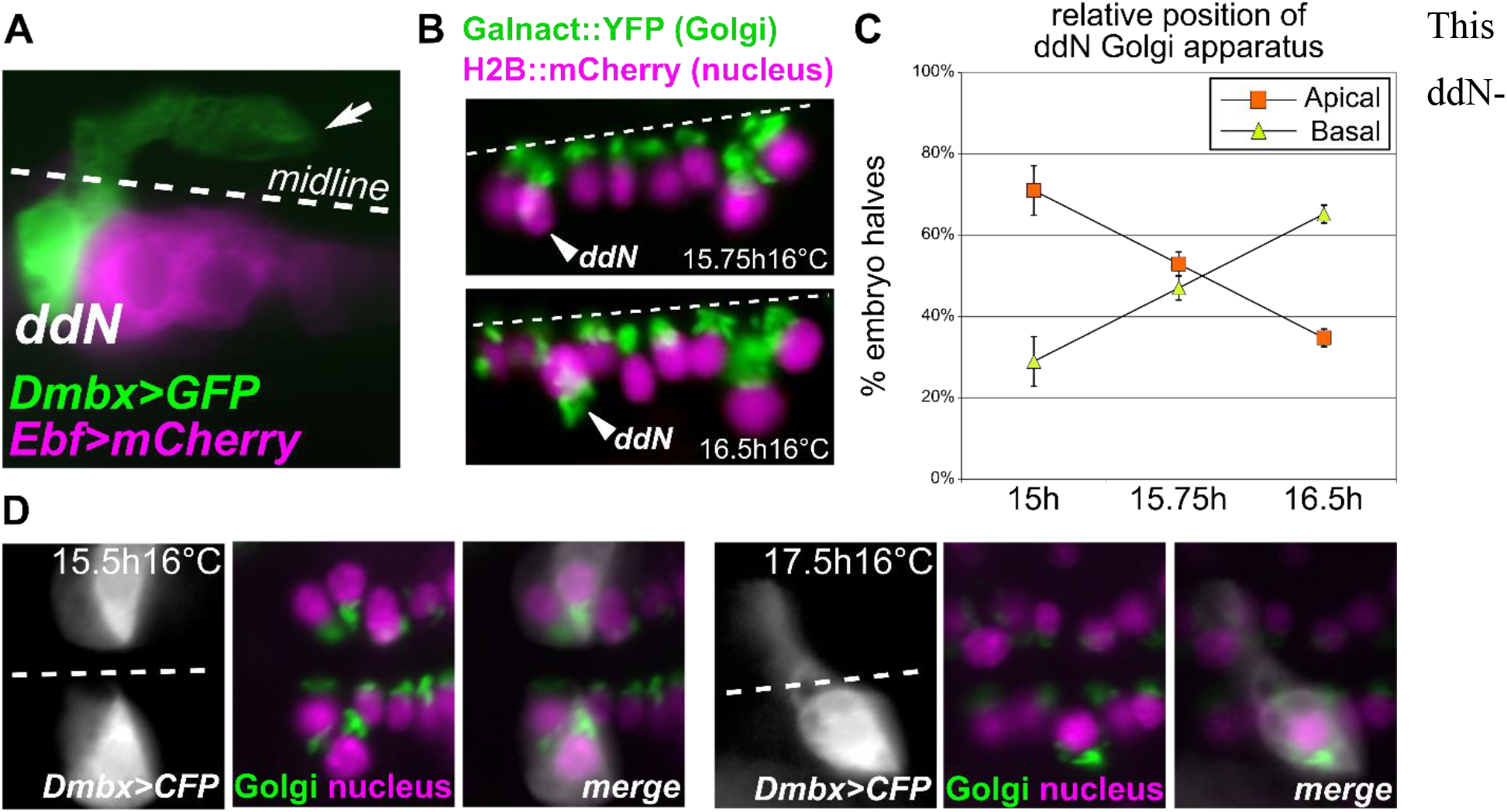
Intracellular polarity dynamics in ddNs during contralateral axon outgrowth. **A)** MG cells labeled with *Ebf>Unc-76::mCherry* and ddN labeled with *Dmbx>Unc-76::GFP*, on one side of mosaic transgenic embryo. Arrow indicated ddN axon growth cone extending posteriorly. **B)** A9.30 lineage cell Golgi apparatuses labeled by *Fgf8/17/18>Galnact::YFP* and nuclei labeled by *Fgf8/17/18>H2B::mCherry* showing intracellular polarity inversion in ddN between 15.75 and 16.5 hours post-fertilization at 16°C. **C)** Plot showing inversion of Golgi apparatus position in the ddNs, showing a shift from a more medial, apical position to a more lateral, basal position relative to cell nuclei. Each time point was analyzed in three independent replicates. In each replicate, 31<n<100. Full data set contained in Supplemental Table 4. **D)** Three-color labeling showing Golgi apparatuses, nuclei, and cell body before (15.5h, left panels) and after (17.5h, right panels) inversion of polarity and axon extension across the midline (dashed lines).

The relationship between centrosome/Golgi apparatus position and site of axonogenesis has been subject to long-running debates. Studies on cells *in vitro* suggested that the centrosome is positioned proximal to the site of axonogenesis (de Anda et al., 2005). However, more recent evidence suggests that *in vivo*, and depending on neuron type, centrosome position does not determine axon outgrowth and can even be *distal* to the site of axonogenesis (i.e. on the opposite side of the nucleus)(de Anda et al., 2010; Distel et al., 2010; Stolfi et al., 2015; Zolessi et al., 2006). Since centrosome repositioning has been shown to depend on microtubule stabilization (Pitaval et al., 2017) the repositioning of ddN centrosomes that we observe might be effected in part by microtubule stabilization, driven by ddN-specific upregulation of *Saxo* and/or *Nckap5* (see follow up experiments below). *Saxo* transcripts were also detected in migrating BTNs by ISH (Fig 3A), and by single-cell RNAseq analysis (Horie et al., 2018a), hinting at the possible involvement of Saxo in centrosome repositioning in BTNs too.

### ddNs upregulate extracellular matrix proteins and the axon guidance cue Netrin1

The extracellular matrix (ECM) has also been shown to drive centrosome repositioning and axon outgrowth (Polleux and Snider, 2010; Randlett et al., 2011). For instance, Golgi apparatus/centrosome position can be precisely manipulated *in vitro* by seeding cultured cells on microprinted Fibronectin-based ECM (Théry et al., 2006). We detected enrichment of transcripts from a gene encoding a relatively short, cysteine-rich predicted extracellular protein whose closest BLAST hits were the N-terminal heparin-binding and collagen-binding domains of Fibronectin-like proteins from various organisms. We termed this gene *Fibronectin-related* (Fn-*related*, *KH.C2.667*, LogFC = 1.7). Expression of *Fn-related* was detected by ISH in the anterior lateral rows of the MG including the ddN and its sister and/or cousin cells (Fig 5A). It was also strongly expressed in the notochord down the length of the tail (Fig 5A **inset**). This pattern of expression is notable because the notochord ECM sheath is composed of various proteins (Wei et al., 2017) including conventional Fibronectin (Kugler et al., 2008; Segade et al., 2016), and could serve as a substrate for axon extension into the tail given its position just ventral to the neural tube. Fibronectin is required for assembly of Fibrillin microfibrils in the ECM, and the region of Fibronectin that interacts with Fibrillin was mapped to the collagen-binding domain (Pankov and Yamada, 2002; Sabatier et al., 2009), which is partially conserved in Fn-related (see **Supplemental Sequences**). In *Ciona*, Fibrillins are encoded by several paralogous genes including *KH.C1.184*, which was also enriched in ddNs by microarray but unconfirmed by ISH (see Table 2). Nonetheless, the expression of *Fn-related* by the ddN itself hints at the possibility that the ddN may be assembling a narrow ECM path for its own axon in the interior of the neural tube, which could facilitate subsequent axon growth towards and across the midline.

**Figure 5.**
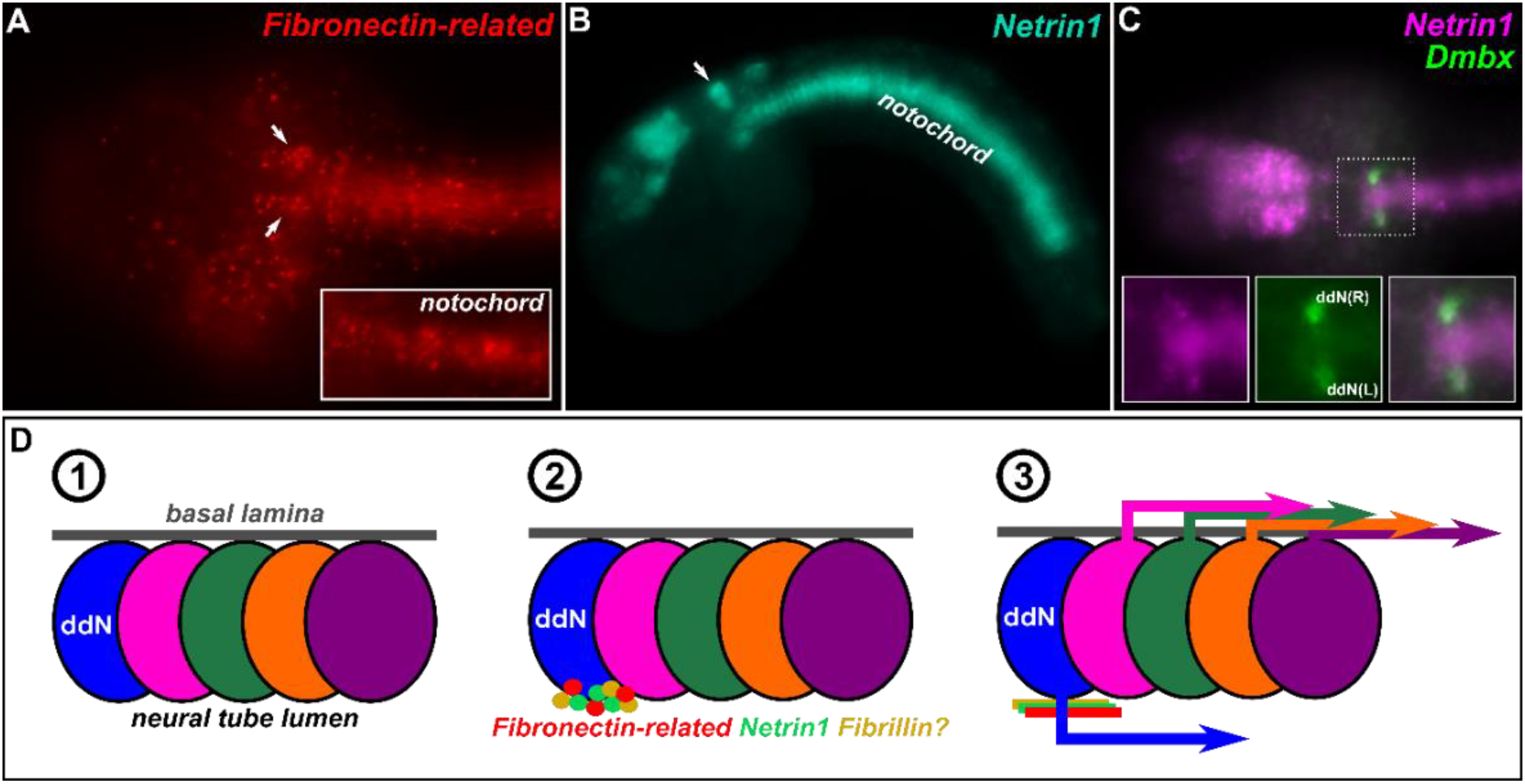
Extracellular proteins expressed by ddNs and model for autocrine mechanism of ddN polarization. **A)** *In situ* hybridization of ECM protein-encoding *Fibronectin-related*, showing expression in ddNs (arrows) and notochord (inset = different focal plane). **B)** *In situ* hybridization of axon guidance cue-encoding *Netrin1*, showing expression in ddN (arrow) and notochord. **C)** Two-color *in situ* hybridization showing co-expression of *Netrin1* (magenta) and known ddN marker *Dmbx* (green). **D)** Cartoon diagram describing proposed model for an *intrinsic* program for ddN polarization, based on autocrine deposition of ECM molecules (Fibronectin-related, Fibrillin) and an axon guidance cue (Netrin1). Briefly: 1) MG neural precursors on one side of the embryo depicted with their basal side pointing laterally, attached to the basal lamina of the neural tube, and their apical side pointing medially, exposed to the lumen of the neural tube. 2) the ddN expresses and apically secretes extracellular proteins including Fibronectin-related, Netrin1, and possibly Fibrillin. 3) This forms an attractive substrate for ddN repolarization and axon outgrowth medially, away from basal lamina and towards the midline, eventually crossing the midline. All other neurons extend their axons laterally along the basal lamina of the neural tube.

We also found that the major axon guidance molecule-coding gene *Netrin1* (*KH.C12.72*, LogFC = 1.3)(Boyer and Gupton, 2018) is enriched in the ddN. We confirmed, by ISH, *Netrin1* expression specifically in the ddNs (Fig 5B,C). Like *Fn-related, Netrin1* is also highly expressed in the notochord, as is another axon guidance molecule, *Sema3a (Kugler et al., 2008)*, further supporting the potential role of the notochord in guiding MG axons into the tail. Although Netrin was long thought to be exclusively a long-range cue (Kennedy et al., 1994; Serafini et al., 1994), tissue-specific targeting of *Netrin1* in the vertebrate hindbrain and spinal cord recently revealed that the role of netrin1 protein in guiding midline crossing is consistent with its function as a short-range cue. More specifically, netrin1 from the floorplate of the developing hindbrain is dispensable for midline crossing (Dominici et al., 2017; Yamauchi et al., 2017), which is mostly regulated by netrin1 distributed along the axon path and derived from ventricular zone progenitor progenitors instead. In the spinal cord, both netrin from both sources act synergistically as both short- and long-range cues to guide midline crossing (Dominici et al., 2017; Moreno-Bravo et al., 2019; Varadarajan et al., 2017; Wu et al., 2019). Furthermore, a role for UNC-6/netrin in *C. elegans* is instructive for neuronal polarization and defines the site of axonogensis *(Adler et al., 2006)*. These short-range functions might explain the potential of ddN-deposited Netrin1 to specify the nascent axon medially and to serve as a short-range cue to drive axon extension towards the neural tube lumen and across the midline, in combination with a suitable ECM substrate potentially formed by Fn-related and Fibrillin (model proposed in Fig 5D). Because Netrin1 might require heparin sulfates (Kappler et al., 2000) as well as a Fibronectin substrate for axon attraction (as opposed to repulsion on Laminin)(Höpker et al., 1999), and Fn-related comprises both heparin- and putative Fibrillin-binding domains (see **Supplemental Sequences**), there is the possibility that Fn-related, Fibrillin and Netrin1 secreted by the ddN itself combine to form a larger complex that couples ECM composition to axon attraction.

Among other poorly studied genes or genes without any obvious or specific function in establishing ddN-specific traits that were confirmed by ISH were *Fam167a* (*KH.C2.629*, LogFC = 3.1), *Calmodulin1-related (KH.C8.573*, LogFC = 2.3) and *Mitochondrial pyruvate carrier* (*KH.C1.85*, LogFC = 1.3)(**Supplemental Figure 2A-C**). Additionally, we could not detect with any certainty the expression of two candidate genes in the ddNs by ISH, *Myosin10* and *Fibrillin* (**Supplemental Figure 2D,E**). These negative results may have been due to poor probe design, which were prepared from short synthetic sequences (~500 bp). However, they also represent potentially false positives in the differential expression dataset, suggesting caution in interpreting such analysis devoid of any confirmatory ISH data.

### MGIN2-enriched transcripts

While several ddN-specific markers have been previously described, there are relatively few known markers of MGIN2 other than *Vsx.* Although *Vsx* was indeed the gene that was most enriched in MGIN2s in our dataset (Table 2, Supplemental Table 2), this enrichment could be attributed to expression of the *Vsx>GFP* plasmid used to sort these cells, which contains portions of the *Vsx* coding sequence.

The second-most enriched transcript we identified in MGIN2 was *Chrnb* (LogFC = 3.7), encoding a beta (non-alpha) subunit of the neuronal nicotinic acetylcholine receptor. *Chrnb* was not found in the current KyotoHoya genome assembly, but is represented by previous gene models. We designed an ISH probe based on the *KYOTOGRAIL.2005.771.2.1* gene model, which revealed highly specific expression in differentiating MGIN2s (Fig 6A). According to the *C. intestinalis* connectome, MGIN2 receives synaptic inputs primarily from Photoreceptor Relay Neurons (prRNs), which in turn receive inputs primarily from Group I photoreceptors (Kourakis et al., 2019; Ryan et al., 2016). Recent experimental evidence suggests that the negative phototactic behavior of *Ciona* larvae is mediated by directional light detected Group I photoreceptors, while Group II photoreceptors mediate a light dimming “shadow response”. In the most recent model of the negative phototactic neural circuit, acetylcholine neurotransmission from prRNs onto the MG might provide the synaptic link between visual processing and motor control (Kourakis et al., 2019). Thus, acetylcholine receptors formed in part by Chrnb subunits might be mediating this crucial step in the visuomotor pathway of *Ciona*.

**Figure 6.**
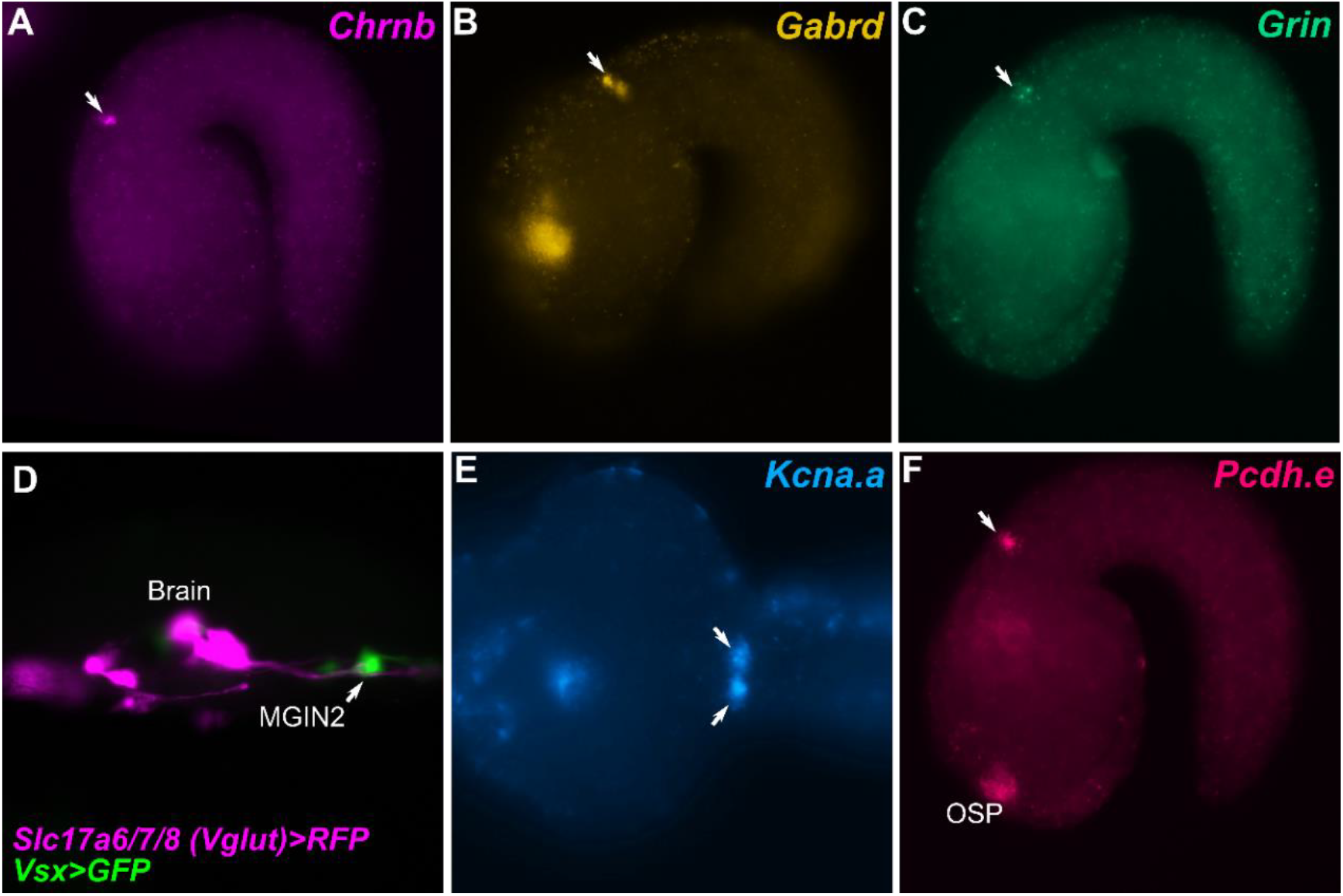
Candidate effector genes preferentially expressed in MGIN2 vs. ddN. *In situ* hybridization of neurotransmitter receptor subunit-encoding transcripts **A)** *Chrnb (neuronal nicotinic acetylcholine receptor, beta/non-alpha subunit)*, **B)** *Gabrd (GABA receptor subunit delta)*, **C)** *Grin (NMDA-type ionotropic glutamate receptor).* **D)** Larva electroporated with *Slc17a6/7/8(Vglut)>RFP* (magenta) labeling glutamatergic neurons and *Vsx>GFP* (green) labeling MGIN1 and MGIN2. Axons from unidentified glutamatergic brain neurons extend and contact MGIN2, suggesting an unknown glutamatergic sensory relay input into the MG via MGIN2. **E)** *In situ* hybridization of transcripts encoding the Shaker-type voltage-gated potassium channel (*Kcna.a*). **F)** *In situ* hybridization of *Protocadherin.e.* OSP: oral siphon primordium. Arrows in all panels indicate MGIN2.

In addition to cholinergic transmission, GABAergic transmission has been proposed to play a minor role in the negative phototactic pathway, and a major role in the shadow response pathway (Brown et al., 2005; Kourakis et al., 2019). We detected enrichment of transcripts from the GABA receptor subunit delta-encoding gene *Gabrd* (*KH.C1.1254*, LogFC = 2.6) in MGIN2, which we confirmed by ISH (Fig 6B). In the future it will be interesting to ascertain whether this localized expression confirms the requirement of GABAergic neurotransmission in negative phototaxis or whether it implicates a cryptic role for MGIN2 in the shadow response.

In contrast to acetylcholine and GABA, no role for direct glutamate neurotransmission onto the MG has been proposed. However, we identified MGIN2-specific enrichment of transcripts for the NMDA-type ionotropic glutamate receptor-encoding gene *Grin* (*KH.S2302.1*, LogFC = 2.2, Fig 6C). While most prRNs that provide synaptic input onto MGIN2 are predicted to be cholinergic and/or GABAergic, we detected the presence of putative glutamatergic brain neurons projecting to and making contact onto MGIN2 in larvae co-electroporated with *Slc17a6/7/8(Vglut)>tagRFP (Horie et al., 2008; Stolfi et al., 2015)* and *Vsx>GFP* reporter plasmids (Fig 6D). The identity of these MGIN2-contacting brain neurons remains elusive. According to the connectome (Ryan et al., 2016), MGIN2 also receives substantial input from other classes of brain neurons, including antenna relay neurons, which presumably relay positional information from the otolith-attached antenna cells, and coronet relay neurons, which presumably relay information of unknown nature from dopaminergic coronet cells. Larval swimming and attachment are modulated by gravity, which is lost in mutants lacking an otolith. Furthermore, the shadow response can be altered by pharmacological treatments predicted to interfere with dopamine that is presumably released by the coronet cells (Razy-Krajka et al., 2012). Therefore, it is possible that *Grin* expression in MGIN2 is necessary for the modulation of swimming behavior by these other pathways, through those observed yet so far unidentified glutamatergic relay neurons in the brain.

Other potential effectors of MGIN2 function that we detected as highly enriched in MGIN2 and validated by ISH included *Kcna.a* (*KH.C1.232*, LogFC = 3.1, Fig 6E), encoding a Shaker-related, voltage-gated potassium channel, closely related to TuKv1 (Ono et al., 1999) from *Halocynthia roretzi*, a distantly related tunicate species. Another candidate was *Protocadherin.e* (*Pcdh.e, KH.C9.518*, LogFC = 2.7, Fig 6F). Protocadherins play numerous roles in morphogenesis, including being extensively implicated in dendrite morphogenesis and dendritic arborization (Keeler et al., 2015). Thus, its specific expression in MGIN2 could be related to its relatively elaborate dendrites, a morphological hallmark of MGIN2 (Stolfi and Levine, 2011)(Fig 1B). We also found 2 genes that by ISH seemed to be enriched in both MGIN2 and MN2 (**Supplemental Figure 3A,B**), but not in other MG neurons: *Slc24a4 (KH.L132.15*, LogFC = 2.9) and *Frequenin (KH.C1.1067*, LogFC = 2.0), further suggesting that many effectors might not be strictly neuron subtype-specific, but that specific combinations of unique and shared effectors may precisely delineate MG neuron functions.

### Conclusions

Here we have used molecular perturbations, embryo dissociation, and FACS to isolate specific neuronal progenitors in the developing *Ciona* MG, and compared their transcriptomes by microarray. Specifically, we have compared ddN and MGIN2 neurons, which are predicted by the connectome to serve as major conduits for various sensory modalities to modulate the larva’s swimming and escape response behaviors. Our transcriptome profiling points to possible effectors of ddN/MGIN2-specific electrophysiological properties (e.g. Scna.a, Kcna.a), morphology (e.g. Saxo, Nckap5, Efcab6-related, Pcdh.e), and functional connectivity (e.g. Chrnb, Gabrd, Grin). These now comprise an attractive set of targets for tissue-specific CRISPR/Cas9 somatic knockouts (Gandhi et al., 2017), in future functional studies of the gene regulatory networks regulating neurodevelopmental processes in the *Ciona* larva.

## Supporting information

Supplemental Sequences

## Acknowledgments

We thank Florian Razy-Krajka for discussions and feedback on data analysis and manuscript. We thank the constant support of Michael Levine and Lionel Christiaen. This work was funded by NIH R00 award HD084814 to A.S., an NSF Graduate Research Fellowship to C.J.J., and a PURA award to J.O. from Georgia Tech.

