## Supplemental Figures 1-3 for "Neuron subtype-specific effector gene expression in the Motor Ganglion of Ciona"

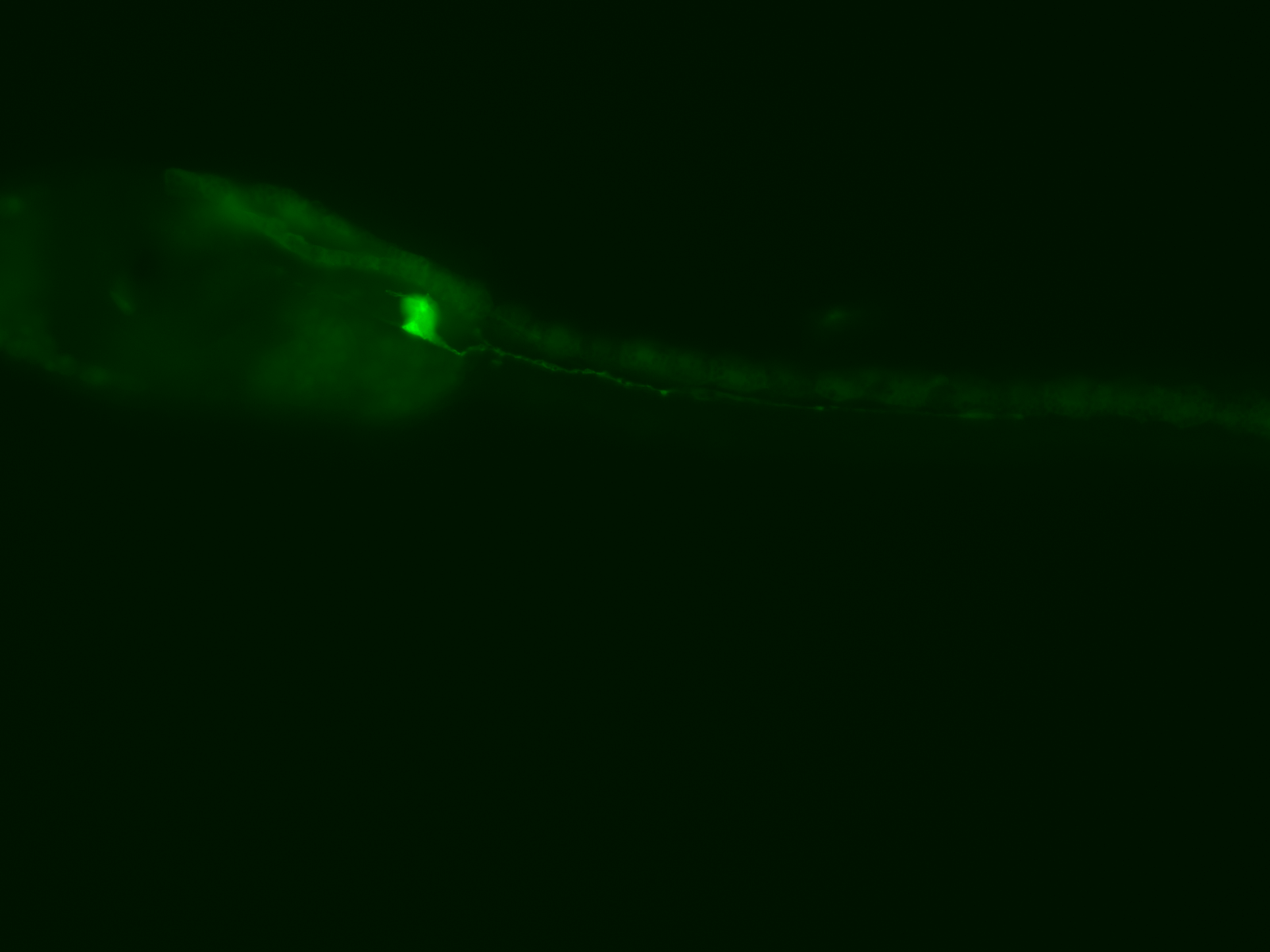

**Supplemental Figure 1.**

Weak GFP expression observed in dorsal epidermis midline in larva electroporated with *Vsx>Unc-76::GFP,* as the likely source of contaminating cells in MGIN2 samples.

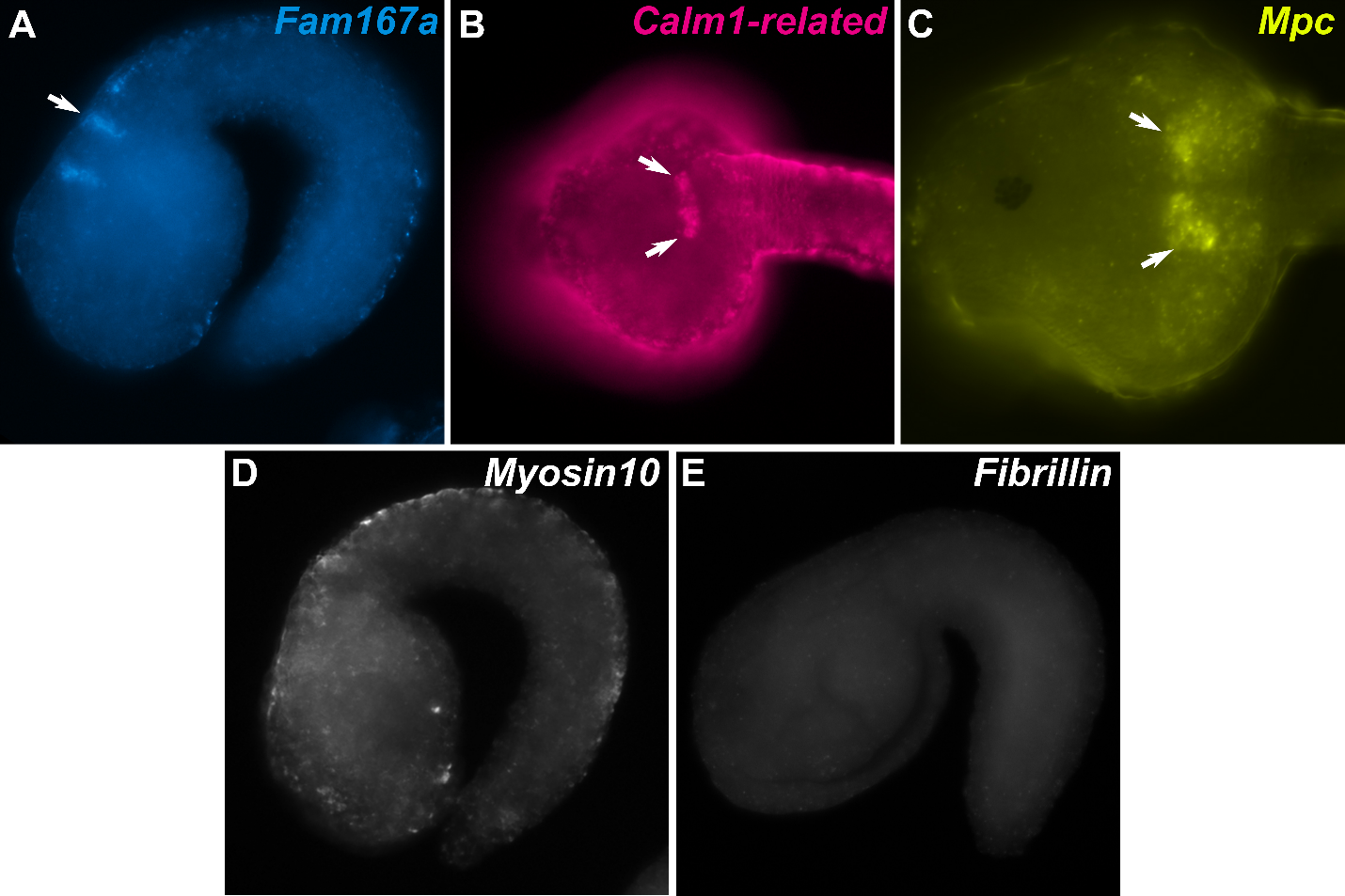

**Supplemental Figure 2.** ***In situ* hybridization for other candidate ddN effectors*.***

Arrow indicates putative expression in ddNs

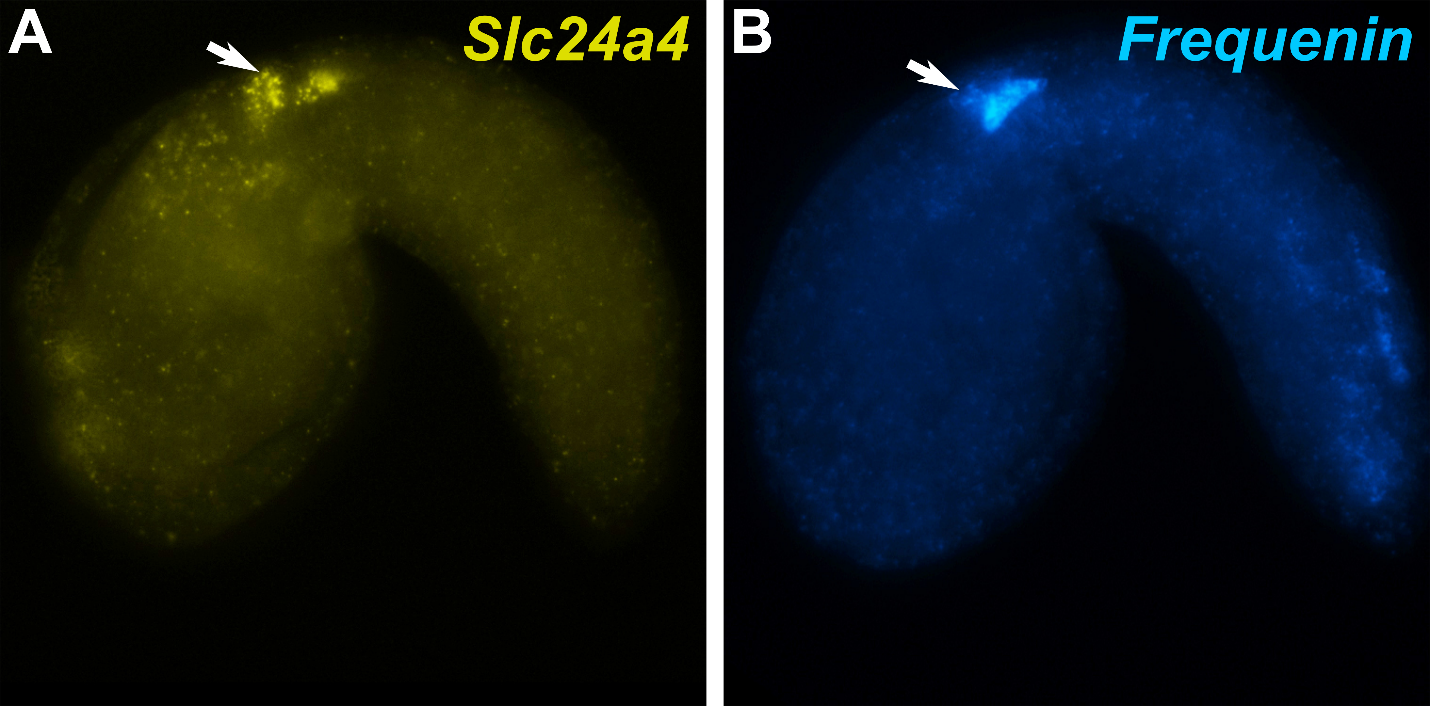

**Supplemental Figure 3.** ***In situ* hybridization for *Slc24a4* and *Frequenin.***

Arrow indicates putative expression in MGIN2.
