## Supplemental Sequences for "Neuron subtype-specific effector gene expression in the Motor Ganglion of Ciona"

**Supplemental Sequences – Gibboney et al. 2019**

**Efcab6-related::GFP coding sequence:**

**Efcab6-related**

**eGFP**

ATGTCGCAAACCATACTCAGTAGACCTGCCAGCCAAGCCAGCAGGGTGGGCATGTTACCCCGCATAGAACACCCCTTATCCCGCCTTGAAAACCCTGGAAACATGTCGGTACGAGGGGTAACAAGGGGCAGCAGTCGGGCGGGTGATGTCCCAAAGCCCCACCGTTTATCACGTGACGCTATGAGCCCCAGACGAAAAGATTTAGACCCGAAGATGCCGATTGATTCGATTCCTGAGGGCATTGAAGTCGTTCACTCAAAAGATCCGAATAAACCGACGTTACCTGTGTTCGGTAATCGAGCTGTTATGAGCCGAGCGGAGTCACGTGGTAGCAACGTCAGTAAAATGACGTCACTGTCAAGACCTGGAACACAAACGCGAATTGAAATTGATGAACTTGAAATGTTGCTACGCGACAAAATCAAGACCGGTGGTTTCTTCAGTGTAAGACAAGCATTCAAAAATAACGACCCAGAAGGTAAAGGAAACGTCACAAAGGAGGCACTCATGCATATCTTGACGTCATTGTTGGGTAGAGTTTTGTCTTCAAAACAATTCCATCTTTTACTAAAACGACTTCACCTTGATGATCGAACTGTCGTCAAATTCGAGGAGTTTTACGCTCAGTTTCGTGAAAGTGTTTCATCTGAATATCCTCGTTGGTTAGACCCGGTTGCGAGGACGCAGACGGAGCGTGCGAGCATGACAGCGTCACAGGTTCACGCCCAGCTAAAGGAGAGAGCCAAACAGAGGTTTCTGGATCTTGCTGATCTTGTTCCACAGATGAATCCAGGAGGCTCTGAGAGGATCATGGCTCCGGAGTTTAGGAACGTTCTTAATAGACTCGGATTCTTCATGAGCGATGCAGAATTTGATAAAATTTGGAAAAAATACGACACATCAAACCTTGGTGTAGTGAAAGCTAGTTCATTGCTCAAAAAACTTGGAATCGAATTCCGAGAATCAAAATCGCCGTCAGCTTCTGTGGCGAGGTCTACAGGAAGTAATAAGAAGGCACAAGTTGTTACTTACGACTCTGCACCTCCAACCGGGCGTTCCATGAGACCCAGTGAAGCGGAACGACAAACATCCATAGATATCGAGAAATGGTTAAAAGATAAATTTCGTGAAGGATTTAAATCAATGCGGAAAGAGTTCAAGAAAGGGGATTTGAAAAATACAGGAAAAGTACCACGTGATGTTTTCCGTCGAGTAATCTCGCAATTTGATCTTTTCCTCCGGGAAGATTCATCTTTAAACATGTTCCTTGCAAGATGTGGATTACCAGAGCATGGTGAGATCTCTTACGTCGATTTCTTGAATAAGTTCCAAGACAGAAGCGCGACAGGGATGGCTCATAATATTCTATCTAACCCAAAACACAGATTCAACAAAGAGCCTCGTCCCATTAGCCCCAAATCCACTGTTACAGCCGTCGAGTCGAAAATGATGACATTATTTCAATCAGACTTCCTTGCTTTGCTTGGAATGTTTCATAAGATCGACAAACATCAAGAAGATGTGATTTCTCAACAAGAGTTCAGAGCTGCCATTGAGAGCAGATTCCAGCTTGAAATGAATGATGATGAATTCAGTCACTTCCTTGAACAAGTTCCATTAAATGAAGATGGTGCAGTCAAGTATCCAGAGTTCATGGCTCAGTTTGATACAAAACAAGGTGCAAAGAGTTTATGGGATGGCAAGTCTGTTGTTTCAAAAGCTGCTGTGTTTGTTCCTACATCTGATAAACCAAAGGAAAGAACTGTTGATGACCTTCATTCAATATTGAGGGTGATTGTCCGTAACGACATGGCAAAGCTGGAAGGAGAATTCAGGCAGCTGGATGAATACAATAGTGGGAAGCTTACACAAGAAATGATGTTCCAACTATTATCAAAAATGCACATTGTACCGTCAGTAACACGAGGTGAAATAAGAAGATTGTGGGAAACTTTCATTGTCAACAAAAACCGAACATTCAGTTTTCTACAATTTGTTCGACATTACGGTTACTCACTCAAATCTGCAGCATTCCCGAATGCCAAGATAGCTCCACCACAACGAGGGGATAATGACTTCATGATTAGGTCAAGGAAACTTAACTGTGCGGCAGATATGTTAGAAGATAGTCTAAGAGCTAAGGTTGATTATTTATGGGAAGACCTTCGTCGTGAGTTTGTTGAGATGGATCCTTACCACACAGGGTTTGTATCACGGGATGAGTTCCGTGATGTGCTCATGGAGCTGTGTGTTCATCTCACGAATCATGAAGCTGAGATCATTTGCAACAAGTTTGAAACAAACACGGATGGAAGAGTTTCATATGTTGAGTTCCTGCGACCATTTGCACAACGTCGGCAGTTATGGAAGGAAGGAAACAACATGCACTCTATACTAACTCACCCACAAGCTGAACTACCTGGATCCATCACTGCAGGGAAACCAACTAAAGGTCTTGAAGCAGTGACTTCTAAACTAAAGCAGCAGCTTGCGGGTGATTGGCGTACTCTAAGAAGGGCGTTTAAAAAGATGGATGTGTCTGCTGATGGAATGCTGACTCTGCCAGAATTTCGAAGCGTTCTTCGACTCTGTAATGTGGTTTTAGATGAAGACGAAGTCTACCATGTTCTTACACAGTATGACAAAGACCTGTCTGGCAAACTGGACTACAAGAAATTCCTGACAGAAAATTTAAGCCGACCATCATCAAAATTATCGGGTGTCtctagtaccatggtgagcaagggcgaggagctgttcaccggggtggtgcccatcctggtcgagctggacggcgacgtaaacggccacaagttcagcgtgtccggcgagggcgagggcgatgccacctacggcaagctgaccctgaagttcatctgcaccaccggcaagctgcccgtgccctggcccaccctcgtgaccaccctgacctacggcgtgcagtgcttcagccgctaccccgaccacatgaagcagcacgacttcttcaagtccgccatgcccgaaggctacgtccaggagcgcaccatcttcttcaaggacgacggcaactacaagacccgcgccgaggtgaagttcgagggcgacaccctggtgaaccgcatcgagctgaagggcatcgacttcaaggaggacggcaacatcctggggcacaagctggagtacaactacaacagccacaacgtctatatcatggccgacaagcagaagaacggcatcaaggtgaacttcaagatccgccacaacatcgaggacggcagcgtgcagctcgccgaccactaccagcagaacacccccatcggcgacggccccgtgctgctgcccgacaaccactacctgagcacccagtccgccctgagcaaagaccccaacgagaagcgcgatcacatggtcctgctggagttcgtgaccgccgccgggatcactctcggcatggacgagctgtacaagtaa

**Gene Collection plasmids for probe templates:**

*Lhx1/5: R1CiGC44f09*

*Pou4: R1CiGC32g05*

*Mnx: R1CiGC42o14*

*Ephrin a.d (Efna.d, KH.C3.716): R1CiGC01j20*

*Scna.a (KH.C9.462): R1CiGC46i17*

*Fibronectin-related* (*Fn-related, KH.C2.667): R1CiGC07e23*

*Mitochondrial pyruvate carrier* (*Mpc, KH.C1.85): R1CiGC24j21*

*Kcna.a* (*KH.C1.232): R1CiGC03f21*

**Probe templates from synthetic fragments from Twist Bioscience:**

*>Saxo (KH.C10.475)*

GAGTTACGTCCCGAAATCCATGCGCGAAGTCGAGCGCTACCCAACACCGGATTGGCTGAAACAACAAGAAGATATTTGGAGGAAGACTGGCATGATGACACAAAAGACCAGAGGGATGACCCCCGAAGCTACCGCTGCGTATATTCGAGCTGCAGCATAATAATACTTCTTTTCCACTTTTTCTCAAAAAGAATTTTTGAGATATTTACAATCTTTATTTACGCTATAGGTTTAGACGTGCGCAGGATTGTTTATAGTTAGGCAAGTTTGTTTTCGAAAATCGCTCATCCCATTCTGGGTCGTCAAGCAAAATGCAGTACAGCCTCAAATGAGTGGCGTTTCACAACACTGTTACTGACCAACGTTATAATGTGATGGGCTTAATAATCGTCGCTTGGGTTGACCTGCATTGCTTACCAATCGGGTAATTTGGCTGCTTGTTTGGGTAATCTTAGAAGGCTCCTATTGTTTATGTACTCTGCCCGTTTTGTATAAAAAAA

*>Calmodulin-related (KH.C8.573)*

ATGTCAGCAACGACAAAGCCTGGAACTGCAAAGCCTGAGATTGCGAAGTCTGAAACCATCAAGCGCACTACTGATTCACAAATGGAAGAATTAAGGGAAGCATTTCGATTTTTTGATCGAAACCAGAACGGCAGTATTGAACCTGAAGAATTGGGATCAGTGATGACCTCGTTGGGATATTGCGCTACAGATTCTGAACTAAAAGATATGATCCACGAGGCAGATGTGGATGGGAATGGGAAGATCGACTTCAAGGAGTTTGTCCGGATGATGGAACTTAAGACCAACGAACGTCCAGAGCAAGCAGAGGACGAAGAATTAAGAGAAGCATTCAAGGTTTTCGACCGGGACGGTAACGGTTTAATTAGTCGTGCGGAATTAAGTCAAGTCATGGGAAATCTGGGCGAACAGCTGAGtGAAAAAGATCTCAATGATATGATTTCTGAAGCCGACAAAAACGGTGATGGTCAAATTGACTATGAAGAATTtGTGCAAATGGTGGCGAAGAAGTGA

*>Myosin10 (KH.C3.562)*

TGTTGTGCATATCATCCTTCAAGGCCAGTTATGCAATACCTCAAGTTCCATCTAAAACGAGTGAAAGAGCGTTACCCCGAAACCCCAGGTGGGGTTTACGCAGCTTTTGCTGAGAAATCAATAAATAAACAAACTTCCCGTCGGCGTGAGATGGTTCCCTCTATACCAGAAATATGCGCGGCGCTCGAAAGACGGGATCTTGTTACTGTTATCAAGTGTTACGGTGGTGCAACATGTGATATCTACATTGACTCTTTTACAACTGCTGGACAGGTTGTACAAAAGTTGAGACGTGGTATGCAACTTGAAGGGAATCGAAACACGTTTGCGCTTTTTGAAAAGAAAGGAACTGATGAAAGAGCTTTGGAACCTGCTACCTTCCTATGTGATAGTGTGGCTAAATTTGAATCACTAAACCACGAGCGAGATATGGCAAGTGGGGAGTTGGAGTGGGAACTTTATTTCAAGCTGTATTGTGTGTTTGATCCCATGGAGGTCAA

*>Fibrillin (KH.C1.184)*

ATACATCAATGAATGCTTGCTACTAAACCATGGTTGCCACACCAAAGCAACCTGCTATAACTTAGATGGAGATTATGTTTGCGAGTGTAATGGTGGTTACAAGGGGAATGGAACATACTGCGAAAATATAGATGAATGTTTAGAGAACACAGCTTACTGCCATAGAGATGCAACTTGTAGCGATACTGAGGGTTTCTATGCTTGTATTTGTAAGCAAGGCTACACTGGTGATGGCTTATATTGTACAGACTTGAACGAATGCAAAGATCCAAACAGCTGCTCTGCCGTAGGATCCGAATGCACCAACCTACCAGGAAGTTACTCTTGCGCTTGTAAACAAGGATACTCTGGGGACGGAAGCCAATGTTCAAAAAAATCCCCACCAAGACCGGCGGACCAGTTCAGTGCAGAAGCTATCGGATCACAAACCAAAGCACAAAGTGCTTCTACAATGCTCGCTCTCTTTGCAACAGTGGGGGCGGTTGTGTTCTTTATGGTTA

*>Chrnb (KYOTOGRAIL.2005.771.2.1)*

GGAAAACACCGACGGTAACTTCACTGTAAGCCATATGACAAGTGCCCACGTGCGGCATGATGGTTCGGTGGCATGGCAACCACCAGCGTCGCTAAAGAGCTGGTGCTACATTGATGTTACTTATTTCCCTTTCGACGTACAAAACTGCACTATGAAATTTGCTTCGTGGACCTATGGAAACAACACAGTACAACTTGTCCCTATGGAAACCAAGGTTGACGTCACAAACCTTCTTCATAACGGGGAGTGGGACATTGTTGACACTCATGTTGAACATACATCAAGTAGTGATTCTTCACTTTATATAAAAAGGAACACCATCGAAACGATAGAAGTTATGTTTGGTCAGAACGGTATATCATATTGTTTTGTTATGAAACGGTTGCCACTATTCTACATTGTGTTTCTGATCACACCGTGTCTAGGAATTTCGTTCCTTACAGCTCTTGTCTTCTATCTTCCCTCTGACTCCCAGGAGAAGATCACTCTCTGCATCTCAG

*>Gabrd* (*KH.C1.1254)*

ACAGATTTCTACGACTCGTTGTCTTCTTGGTAATAACAGCAACTTTCGGGGCATGCAGTGAAACCGCCACTACAACAAGACACACGAAAACACGGGCACCAGACGAAAAACCTGACCAGGCGTTCTGGGGCGGAGTATTCACGCCCCCGCTGCCACAAGATAGTAGCCGCTCTAATGACAGACATGGAAGCGAAATAACAAAACAGATCGAGGAGATGTTTCGAGACCATAATTATGAGTTGAGCATTCGGCCCAGAAACCATGGAAAACCAATACGTGTTGGTCTGGCCCTGATGATCGAAAGCATTACTGACATTTCGGAAAAGAACATGGACCTGACGTTTACGCTATGCATGCACGAGACCTGGACGGACACTCGGTTAAAATTTATCTCAAATTCTGACGACAACAGCATTGTGCTACCGAGTAGATTAATCAGCAAGCTATGGGTGCCAGATCTGTACATTGTCGGTTCAAAGAGCTCATTTATCCACAAAACT

*>Grin* (*KH.S2302.1)*

TTGAATTTGACTCCATATCTAAACAAGTGACAAGAAATTCTTATGAAAACTCAAGTTCCGCGATTGAAGATCTGAAACTGGGGAAACTGGAAGCTTTTATATGGGATAGTGCTTCATTGCAGTATGAGAGATCGGTTAATTGTGACCTGGTAATGGCTGGAGAACCAACATTTGGACCTGGGTTTGGGATTGCTATGCGCAAGAAGGATAAGTTACTGGATCAAATTTCATTGCTTCTGCTGAGTTTTCATGAGAATGGGTTCATGGACAGTTTGAGGTTGAAGTGGACATTGAACCAAGGCTGTCCCCAACGTACATCAAGTCCCGCAACCCTCAAGCTCGAAAACATGGTCGGGGTTTTTATATTGATCGGTATAGGGATTATTTCTGGCGCTATTTTCACAGCTATTGAGGTGGTTTATAAAAGAAAGAGGAATTAGTTGAGCGTGAATAGGTGTGAATATGTGGGGTTTTGTGTTATTGTAGTAGATATAGTGGAA

*>Protocadherin.e* (*Pcdh.e, KH.C9.518)*

TAAGTGCACTGAGTTGTGTAAAACATATGGACACTGTGATACGTGCTGGATGCCAAGCTCAGGTTATCCCGTGGAGCAGACGAGCCAAATTCCAAATGACGTCACAACACCATGGCAACCGAAGTTCTCCGCCACGAGGAAGGATAGCGGATGTTCTGTCGAAACTATGTCGTCCGCGCTCTCTTATTACAAGCTGACGCGGTCTGAGCAACAGCGACGTCGTAATGATGACACACTTCCAGACCTTGTAGCGAAGGGGCGTACAGCCATGGGCAGTGACGCTAATTCTTTGATGTCTTCAACAAACTCTTCAGGAGCGTCATCGACACGCACCGGTGACCTTAAACATACGTCATCAATTCAAAAATCGCCAACTCATCGGGCTTTGAAAAAGTCCAACAGCGATATAGGGTATGAAGCGAAAGACGCAACACCACGGCAGGTCGTTATTCCTAATAGCCTATCCACACAATGCTAAAGTCGCCTCCCAGGTTTTCCAT

*>Slc24a4 (KH.L132.15)*

CCCCACAAACAGAAGATCAAGTTGATGATGGGTTGATATGTGATGAATGTTTGCAAGCTCAGCAATACGAGACGAGATACAAAGAGGTTGGTGAAGGAGTTGATTTGGGAGGTTGCGATAAAGAAGAGGGAAGAGATGGATGTTGCCGACATCAAAAGAAGAGTCCGCCTCCTCCTTATTCAGAAAATGAAGAAAATTTCAGCAAAGAATTTGAAGCTGGATTGAGCGACAACACAGTCCAACCCTCAAGAAACTCGAACCAAAGCAATCATAATCGATTCATTTTCCACCCAAGGAGTTGTTCGTGCGGACATTTTGGGACAAACCCAGACTTTTCATCAGTTGACAACGATGTCGACGCTTACAGTGAAGTAGCGAGCGTGGGTGATGTCATGTCTGGGAGACCTCAAGTGCCAGATGAACCTGAGTCACCTGACCAACCAACATCAGAGGACAACCCTCCTGTCGCAGGTGAAGCAGGATTGAATGAGAATGTTGGA

*>Frequenin (KH.C1.1067)*

AAGGAGGAATTTCAAAAGGTTTACCAGCAGTTCTTTCCCAAAGGAAATCCGTCAAAATTTGCAAATTTCGTTTTTAACGTTTTCGACTCTGACAAGGACGGCTTCATTACGTTCAAAGAATTTATTTCTGCCCTTTCCGTGACGTCACGAGGAAACCTGGACGAAAAGTTAGATTGGGCTTTCAATCTTTACGATCTTGATCATGACGGTTTTATTACAAGAGAAGAGATGTTAAACATAGTAGATGCCATCTACTCCATGGTGGGAAACGCTATGGACTTGCCGGAGGATGAGAACACACCTGAGAAGAGGGTTAACAAAATATTCTGTCAGATGGACCAGAACAAGGATGGCAAATTGACGAAAGATGAATTCCGAGAAGGCTCGAAATGCGATCCCTATATCGTGAAGGCGTTGTCTGCTGGACTAGGAGGGGCGGAAAGCTGCCCGTCGTGAATTGATTTTAGAAAGTTTGCCAATGTAATTAAGCATGTTACCCA

**Probe templates from *de novo* cDNA:**

*>Dmbx*

GCAATTCGTGCAATGTCAGTGTTCAACGCATTACAACAACATTACCCAAACACCCAACACCCTTATAACGGCTACAATAGTAACGGGCAAATTGCTGGATCTCTGGCAGATTTAATACTCAAAGGCGGGACTTTCCCCCGCAAACACCGACGCAGCAGAACAGCCTTCACTGCCATGCAACTTGACGCGTTGGAAAGAACGTTCAAAGATGGGCAGTACCCGGACGTAGAAACGCGAGAAAGTTTGGCAATCTGTACAAATTTAGCTGAAGCCAGGATACAGGTTTGGTTTAAAAACCGAAGAGCAAAATATCGAAAACAACAGAGAATGTTGAAAACGCAGGAGTGTGTTCCCGTTACAGATTCGAACGAAACATCCAACAAAGCTGATTCTTTAAACAATGAATCAGCGACAACATCTATGAGAGTGAAAACAGAAAGAACAAGTGATTGTGACGAAACAAAGGACGGTGGTTTCCCAGAAAAAGATACGAAACCAAAAATTGAAAAGAATCCGTTCAGTTTGCCCCCACCTCTAGTTGCTGGTCCATTCATTCAACCAAATATTCATGGAATGTTGGATCTGAGCATCAGGGGATCAAACTACCAGTTGCCGACAACCTCCCCCTACCCTCCCCTTGCCACAAGAATGCTACAAGTTCCTTTCCAACCTTGGAATCTTCAATTTAGCAATCAATCCAAGAAATGATGATGTAATCGTTATATCTGTGATGTCATAATACATGAGCTGGCACTCACCCCATCATGGCTGTTTTATCGATAATTTATTGCACATAACTTTTATTCGTGTCGATTTAATGCGCTGCACACTGT

*>Efcab6-related (KH.C1.1218)*

CGACAGGGATGGCTCATAATATTCTATCTAACCCAAAACACAGATTCAACAAAGAGCCTCGTCCCATTAGCCCCAAATCCACTGTTACAGCCGTCGAGTCGAAAATGATGACATTATTTCAATCAGACTTCCTTGCTTTGCTTGGAATGTTTCATAAGATCGACAAACATCAAGAAGATGTGATTTCTCAACAAGAGTTCAGAGCTGCCATTGAGAGCAGATTCCAGCTTGAAATGAATGATGATGAATTCAGTCACTTCCTTGAACAAGTTCCATTAAATGAAGATGGTGCAGTCAAGTATCCAGAGTTCATGGCTCAGTTTGATACAAAACAAGGTGCAAAGAGTTTATGGGATGGCAAGTCTGTTGTTTCAAAAGCTGCTGTGTTTGTTCCTACATCTGATAAACCAAAGGAAAGAACTGTTGATGACCTTCATTCAATATTGAGGGTGATTGTCCGTAACGACATGGCAAAGCTGGAAGGAGAATTCAGGCAGCTGGATGAATACAATAGTGGGAAGCTTACACAAGAAATGATGTTCCAACTATTATCAAAAATGCACATTGTACCGTCAGTAACACGAGGTGAAATAAGAAGATTGTGGGAAACTTTCATTGTCAACAAAAACCGAACATTCAGTTTTCTACAATTTGTTCGACATTACGGTTACTCACTCAAATCTGCAGCATTCCCGAATGCCAAGATAGCTCCACCACAACGAGGGGATAATGACTTCATGATTAGGTCAAGGAAACTTAACTGTGCGGCAGATATGTTAGAAGATAGTCTAAGAGCTAAGGTTGATTATTTATGGGAAGACCTTCGTCGTGAGTTTGTTGAGATGGATCCTTACCACACAGGGTTTGTATCACGGGATGAGTTCCGTGATGTGCTCATGGAGCTGTGTGTTCATCTCACGAATCATGAAGCTGAGATCATTTGCAACAAGTTTGAAACAAACACGGATGGAAGAGTTTCATATGTTGAGTTCCTGCGACCATT

*>Netrin1* (*KH.C12.72)*

ACCACGGTAGAACGTGGGTACCATACCAATACTATGCTGAAAACTGCCTGAGAAGGTTTCATAAGCCTTATAAAAGAGAAGCGAATGAGACAAATGAACAAGAGGTNCTGTGTTCAGAAGATTTCAACANTTTGNATCCNTTCAGTCTCGGTGCTCTGGTGTTTAACCCGAAAGATGGGCGGCCTTCTGAGGANGATTTCGAACATATTGCTACCCGGGCAAGCAAGACTGAGTTNCGNCATACAGGTGTTCGCGTGNCTGGTGGGACAGATATTTTAGTCAGCAACCCTGGCGCTACTTCCACTCCGTATAGTTTTGGNTCTAGCTCNTATTATGCGATCAATGATATCAGTATTGGAGGTGTTTGCAAGTGCAATGGTCACGCTGATAGATGTATAAAGAAGAAGGGGAAGATGGTTTGTGATTGTAAACACAAGACAACGGGACCGAACTGCGAGACATGCAAAGCTTTNCATAAGGACCGACCGCCTAAGCGAGCTACGAAAACAAACGCGAACGAGTGNATTGCTTGNAATTGCAANATGCACGCCAAGAGATGCCGATTCAGCGCCGACTTATTTGAAAAGTCGGGTGGTACAAGCGGGGGAGTTTGTGTTAAGTGTAGACACAACACACATGGCCGATATTGCCATAAATGTAAGCCTGGATTTTACAGGAACAACAAAACCAAGTCGATGTCGAACAGGAGAGCTTGTAGGCGAATTCCTGTTGTAACTGGCGCNACCACAGCTAAACCAGCTTGCCCACGAACATGTAGCAGCTCAAAAGCTGTAAGAATACCTCCAGCTAAATATTGTAAGATGGAATACGCNGTGATGGTAAGAGTAACCGGGAAGATCAGGATAGGAAGAAGGACCAGCATCACTTACCAAGTACTCNGGGTGTTTAAGAGGGGAGGCATGAGGTTGAAGAAGAAGGAGATGATTGACATCCAGGTCAGGGATCGAGTCCTACAGTGTCTATGTCCCAGGTTGCGAGT

*>Fam167a* (*KH.C2.629)*

TTGTTGCATTAGTTCCAACGTAATATAGCATTTGTAAGTTATATTTCTGTCATTAAAATGTTGGAAACAGAAAACCGCATATCGAGCTACACGAAAAGTTCTGGGTGTTCGAGTGATATTGGTTGTGATGATCTGGCAAAAGTGAAAGCGTTGTCTAAAAAGCTCAAGCTGTCGACCAGGCGCAGGTCTGTACTGCAATGGCAAACCACGATCGGCGAGAAGCATTGTTTGAGAAACTCTGTAACCGAAGACACACTCGAAGCTATCGTGGAACAAACTTCGCCCGTTCATGACGTCACAGAACCGAATTTTGTGACGTCATCGAGCATTTTAGAAAATGATGACGTAACAAATATGGTGAAACTCTGTCAACCAAACACACAAGATGTATCGGTCAATGAAACTAGTACAACAGACTGTATAGAACAGGAAACAACAAAAGATATAAAACTTACGAAACCGGAACAACATGAGAAACTACCTGAACGTTGTAGAAATATCACCGATTCCCTTATCAGCATTCGTAGAGAATTGATGTTGATGCGTATGGAAGATCACACTATTTCACGCAAACTGCTAGACATTCGTGAAGAAATGAAACGAATGCGAGTGAATCAAATGTGTGACGAACATGCCGCCATGTTGGATGACTTCTCATGGGAAATCGACGAAGAAGAACATTTAGATCTAAAACTAAAAGCTGTGAGCGACCTTTCGAGTGCTCTTGTACGAAAACCTGATTCCTTGCTGTGGAGTTCCTCCCCGTTACGTCACATTGGAATCACAAAGAAGTCGATGGAGTTTCGAAGATTTTCTGTCATTTGAAAATATTTTTCCAACTTTTTGTTCTATTTCTTAAAAGTTTTTTTTTGTAGATTGTTCTTTTATTCTTGTTTCGGTATTTCTTTATATTTTGTTGCTGCTGTTTCGTTGTTAAACTAAACGCAATTATAGATTATCGCTTGGAGACGCACGGTTTAT

*>Nckap5 (KH.C9.229)*

TGGTTAAACGGCTCATTTTCATAGAATGAAGCCGAGGGTTGAGAGTCACTACAAGCTGAGGTTTGGTTCAACGTGTCTGAAGAATCGGATTGTAAAACCAAACTCGACGGTCTTTGTGTGGTTCGTTTACCATGAGAGTTTTTGTTTTGAGATGCAGGATTCAATCGATCGTCATTTTTAGTCGAATCCTCTTTGTATCTTCTGAGCTCAACTTCAATGTAGGAATTATTGTCGAAAGACTGACTATTTTCTAAGTTACTTCCGCTTGTGCTTTTTTTAATTTTAAAAAGTTTTCTGTTGTTCGTTTTTTTGCACGATTCTTTTCTCTGTTCCAGGGATTCACCCGAACATGATCTTGAAGGACTCAGTGGAGGCTTGGAGGAGGAGGAATGTCTTTGGGGGCAAGATGAGCAAGACTGCAATGCACATTTGCTGGACGGTGTGGCGTAATCAGCATGTTCGTTCGTTGTATAAGGTTCAGGATGCTCATGTATTATATTAAGTGAACCAACTGATGCATTATCACTGTTTGGGATATTAGGTGTCTTTATTTGTTCCGAATTAACATCGGAAACTGTTGACTTCTGTAAAGCAGAATATAGTTCTACGACTGTAGCAACATCAGTTGCTTTGTAAGGATCAGAATCTTCAGTGTTAGGTGAAGACTGAAGACGTCCCATCGCTTTGGTGAAAATCTTATCTGTTAACCCATTTTTGGGTTTTCTCTCAACGCTGCTGTGCAAATCTCTCATTGGGTCGTTTGAATCTTCTATCAAAACTTGTCTATTGTTTTTACCATGTTCTTTTTCAACTCCGGAAAAAGAAGTTTCATTTTGTTTTTGCAATATTTTGTTATATTTGGAAAGATTGTCAACTTGTCGTTTTAAATTGAAGCACTCAGATTGCAACTGACGTTGAATGAGTTCCGATTGGACCACCTGAGCTGCAACCTTGTTATATTCCACTTTCTGTTTTTCAGCTAGTTTGATAAGCTGATAATTTTGTTCTTCCATTTTTTGAACTCTTTCAGCTAACAAAATGCAGTGTCTTCGAAAATCATCAAGGTCCAGGACTGCTTGCCGCATATCTCTCTGACAATGCATTTGATTGTTATTAGATGTTGAAGTGAAGTTTTGTAATTCCATATTCCTCTTTTGCTCAGTTGCACCAACATATATAGATGGTTTCAATGGTTGAGGCTGTTCTTGTAAAGTTACATAGCGTCCAAGATCTGTGTGATGTTCTGTATGACGTAAAGCGTCACGAAGAGATTCTTCCATGTTGTCAAACTCAGACTTTAATTCCATGGTT

***Human EFCAB6 alignments with Ciona KH.L125.4 (Efcab6) and KH.C1.1218 (Efcab6-related)***

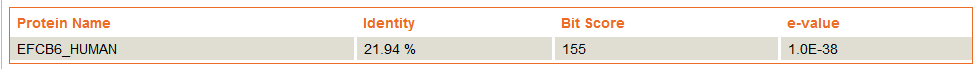

```
CLUSTAL format alignment by MAFFT (v7.408)
Flagged as EF-HAND domain in SMART

EFCAB6          MCKMAIIPDWLRSHPHTRKFTHSRPHSSPCRVYSRNGSPNKFRSSSTTAVANPTLSSLDV
KH.C1.1218      MSQTIL----------------SRPASQASRV----------------------------
                *.:  :                *** *...**

EFCAB6          KRILFQKITDRGDELQKAFQLLDTGQNLTVSKSELRRIITDFLMPLTREQFQDVLAQIPL
KH.C1.1218      ------------------------------------------------------------


EFCAB6          STSGTVPYLAFLSRFGGIDLYINGIKRGGGNEMNCCRTLRELEIQVGEKVFKNIKTVMKA
KH.C1.1218      ---GMLP-----------------------------------------------------
                   * :*

EFCAB6          FELIDVNKTGLVRPQELRRVLETFCMKLRDEEYEKFSKHYNIHKDTAVDYNVFLKNLSIN
KH.C1.1218      ------------------------------------------------------------


EFCAB6          NDLNLRYCMGNQEVSLENQQAKNSKKERLLGSASSEDIWRNYSLDEIERNFCLQLSKSYE
KH.C1.1218      ------------------------------------------------------------


EFCAB6          KVEKALSAGDPCKGGYVSFNYLKIVLDTFVYQIPRRIFIQLMKRFGLKATTKINWKQFLT
KH.C1.1218      -----------------------------------RI-----------------------
                                                   **

EFCAB6          SFHEPQGLQVSSKGPLTKRNSINSRNESHKENIITKLFRHTEDHSASLKKALLIINTKPD
KH.C1.1218      -----------------------------------------------------------E
                                                                           :

EFCAB6          GPITREEFRYILNCMAVKLSDSEFKELMQMLDPGDTGVVNTSMFIDLIEENCRMRKTSPC
KH.C1.1218      HPLSRLE------------------------NPGNMSV----------------------
                 *::* *                        :**: .*

EFCAB6          TDAKTPFLLAWDSVEEIVHDTITRNLQAFYNMLRSYDLGDTGRIGRNNFKKIMHVFCPFL
KH.C1.1218      -------------------RGVTRG-----------------------------------
                                     :**.

EFCAB6          TNAHFIKLCSKIQDIGSGRILYKKLLACIGIDGPPTVSPVLVPKDQLLSEHLQKDEQQQP
KH.C1.1218      ------------------------------------------------------------


EFCAB6          DLSERTKLTEDKTTLTKKMTTEEVIEKFKKCIQQQDPAFKKRFLDFSKEPNGKINVHDFK
KH.C1.1218      ---------------------------------------SSRAGDVPKPHRLSRDAMSPR
                                                       ..*  *..*  . . :. . :

EFCAB6          KVLEDTGMPMDDDQYAL-LTTKIGFEKEGMSYLDFAAGFEDPPMRGPE----TTPPQPPT
KH.C1.1218      RKDLDPKMPIDSIPEGIEVVHSKDPNKPTLPVFGNRAVMSRAESRGSNVSKMTSLSRPGT
                :   *. **:*.   .: :. . . :*  :. :.  * :. .  **.:    *: .:* *

EFCAB6          PSKSYVNSHFITAEECLKLFPRRLKE-SFRDPYSAFFKTDADRDGIINMHDLHRLLLHLL
KH.C1.1218      QTR-------IEIDELEMLLRDKIKTGGFFSVRQAFKNNDPEGKGNVTKEALMHILTSLL
                 ::       *  :*   *:  ::*  .* .  .** :.*.: .* :. . * ::*  **

EFCAB6          LN-LKDDEFERFLGLLGLRLSVTLNFREFQNLCEKRPWRTDEAPQRLIRP--KQKVADSE
KH.C1.1218      GRVLSSKQFHLLLKRLHLDDRTVVKFEEF--YAQFRESVSSEYP-RWLDPVARTQTERAS
                 . *...:*. :*  * *   ..::*.**   .: *   :.* * * : *  : :.  :.

EFCAB6          LACEQAHQYLVTKAKNRWSDLSKNFLETDNEGNGILRRRDIKNALYGFDIPLTPREFEKL
KH.C1.1218      MTASQVHAQLKERAKQRFLDLADLVPQMNPGGSERIMAPEFRNVLNRLGFFMSDAEFDKI
                ::..*.*  *  :**:*: **:. . : :  *.  :   :::*.*  :.: ::  **:*:

EFCAB6          WARYDTEGKGHITYQEFLQKLGINY------SPAVHRPCAED-YFNFMGHFTKP---QQL
KH.C1.1218      WKKYDTSNLGVVKASSLLKKLGIEFRESKSPSASVARSTGSNKKAQVVTYDSAPPTGRSM
                * :***.. * :. ..:*:****::      *.:* *. ..:   :.: : : *   :.:

EFCAB6          QEEMKELQQSTEKAVAARDKLMDRHQDISKAFTKTDQSKTNYISICKMQEVLEECGCSLT
KH.C1.1218      RPSEAERQTSIDIEKWLKDKFREGFKSMRKEFKKGDLKNTGKVPRDVFRRVISQFDLFLR
                : .  * * * :     :**: : .:.: * *.* * .:*. :.   ::.*:.: .  *

EFCAB6          E-GELTHLLNSWGVSRHDNAINYLDFLRAVENSKSTG------AQPKEKEESMPINFATL
KH.C1.1218      EDSSLNMFLARCGLPEH-GEISYVDFLNKFQDRSATGMAHNILSNPKHRFNKEPRPISPK
                * ..*. :*   *:..* . *.*:***. .:: .:**      ::**.: :. *  ::.

EFCAB6          NPQEAVR-KIQEVVESSQLALSTAFSALDKEDTGFVKATEFGQVLKD-FCYKLTDNQYHY
KH.C1.1218      STVTAVESKMMTLFQSDFLALLGMFHKIDKHQEDVISQQEFRAAIESRFQLEMNDDEFSH
                ..  **. *:  :.:*. ***   *  :**.: ..:.  **  .::. *  ::.*::: :

EFCAB6          FLRKLRIHLTPYINWKYFLQNFSCFLEETADEWAEKMPKG-----------PPPTSPKAT
KH.C1.1218      FLEQVPLNEDGAVKYPEFMAQFD------TKQGAKSLWDGKSVVSKAAVFVPTSDKPKER
                **.:: ::    :::  *: :*.      :.: *:.: .*           *.. .**

EFCAB6          ADRDILARLHKAVTSHYHAITQEFENFDTMKTNTISREEFRAICNRR--VQILTDEQFDR
KH.C1.1218      TVDDLHSILRVIVRNDMAKLEGEFRQLDEYNSGKLTQEMMFQLLSKMHIVPSVTRGEIRR
                :  *: : *:  * ..   :  **.::*  ::..:::* :  : .:   *  :*  :: *

EFCAB6          LWNEMPVNAKGRLKYPDFLSRFSSETAATPMATGDSAVAQRGSSVPDVSEGTRSALSLPT
KH.C1.1218      LWETFIVNKNRTFSFLQFVRHYGYSLKSAAFPNAKIAPPQRGDNDFMI------------
                **: : ** :  :.: :*: ::. .  ::.:.... * .***..   :

EFCAB6          QELRPGSKSQSHPCTPASTTVIPGTPPLQNCDPIESRLRKRIQGCWRQLLKECKEKDVAR
KH.C1.1218      -------RSRKLNCA---------------ADMLEDSLRAKVDYLWEDLRREFVEMDPYH
                       :*:.  *:               .* :*. ** :::  *.:* :*  * *  :

EFCAB6          QGDINASDFLALVEKFNLDISKEECQQLIIKYDLKSNGKFAYCDFIQSCVLLLKA-KESS
KH.C1.1218      TGFVSRDEFRDVLMELCVHLTNHEAEIICNKFETNTDGRVSYVEFLRPFAQRRQLWKEGN
                 * :. .:*  :: :: :.:::.*.: :  *:: :::*:.:* :*::. .   :  **..

EFCAB6          LMHRMKIQNAHKMK---EAGAETPSFYSALLRIQPKIVHCWRPMRRTFKSYDEAGTGLLS
KH.C1.1218      NMHSILTHPQAELPGSITAGKPTKGLEAVTSKLKQQLAGDWRTLRRAFKKMDVSADGMLT
                 ** :  :   ::     **  * .: :.  ::: ::.  **.:**:**. * :. *:*:

EFCAB6          VADFRTVLRQYSINLSEEEFFHILEYYDKTLSSKISYNDFLRAFLQ---------
KH.C1.1218      LPEFRSVLRLCNVVLDEDEVYHVLTQYDKDLSGKLDYKKFLTENLSRPSSKLSGV
                :.:**:***  .: *.*:*.:*:*  *** **.*:.*:.**   *.
```

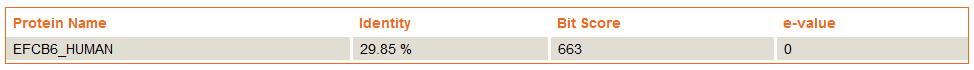

```
CLUSTAL format alignment by MAFFT (v7.408)


EFCAB6          MCKMAIIPDWLRSHPHTRKFTHSRPHS-----SPCRVYSRNGSPNKFRSSSTT-------
KH.L125.4       MSAVQVI-----ERPHPGANGGLRPSSSGVVFSPRPLLSTGRNVRSLRNSTTSLARSFHV
                *. : :*     .:**.      ** *     **  : * . . ..:*.*:*:

EFCAB6          -----------AVANPTLSSLDVKRILFQKITDRGDELQKAFQLLDTGQNLTVSKSELRR
KH.L125.4       PSTSSGSTTVMLAANPKLTSIEVEAIIKQKLWENMEPLKQAFQLYDADRCTNVTKGEFRR
                            .***.*:*::*: *: **: :. : *::**** *:.:  .*:*.*:**

EFCAB6          IITDFLMPLTREQFQDVLAQIPLSTSGTVPYLAFLSRFGGIDLYIN-GIKRGGGNEMNCC
KH.L125.4       VLENYCLPMTAEQFNALVVKIGTNPNGTISYLRFIDKFVTNSLTPNIGVARILSRPSSSN
                :: :: :*:* ***: ::.:*  ...**:.** *:.:*   .*  * *: *  ..  ..

EFCAB6          RT-LRELEIQVGEKVFKNIKTVMKAFELIDVNKTGLVRPQELRRVLETFCMKLRDEEYEK
KH.L125.4       KQRIDTIERFLQQKIAANLKNVIRSFKLFDYNMDGLVQRHELRRVIENYCLKLNDSQFNK
                :  :  :*  : :*:  *:*.*:::*:*:* *  ***: :*****:*.:*:**.*.:::*

EFCAB6          FSKHYNIHKDTAVDYNVFLKNLSINNDLNLRYCMGNQEVSLENQQAKNSKKERLLGSASS
KH.L125.4       LWSRYDFQHTGAINYRDFLRRLGVNAATAERHVTTPLNNGPDEER---DDEDGGCGERVS
                : .:*::::  *::*. **:.*.:*     *:     : . ::::   ..::   *.  *

EFCAB6          EDIWRNYSLDEIERNFCLQL-------SKSYEKVEKALSAGDPCKGGYVSFNYLKIVLDT
KH.L125.4       EEGWEGWVVKGYNIPLVIKLLTYIQDMKNNYQNIKRALMTFDISSDGFISIDDLKAVLDN
                *: *..: :.  :  : ::*       .:.*:::::** : * ...*::*:: ** ***.

EFCAB6          FVYQIPRRIFIQLMKRFGLKATTKINWKQFLTSFHEPQGLQVSSKGPLTKRNSIN----S
KH.L125.4       FVLPTSDEIFHQLMYKFEVRGTGKVSWEQFLSKFQDPQSNGNGQTIPIGQNHKVNPTRQA
                **   . .** *** :* ::.* *:.*:***:.*::**.   ... *: :.:.:*    :

EFCAB6          RNESHKENIITKLFRHTEDHSASLKKALLIINTKPDGPITREEFRYILNCMAVKLSDSEF
KH.L125.4       LDNSSTPDVIKKLKDHILNNFATLKQAFLAFDENRKGVIGRKDLRRIIETFAMKISDENF
                 ::* . ::*.**  *  :: *:**:*:* :: : .* * *:::* *:: :*:*:**.:*

EFCAB6          KELMQMLDPGDTGVVNTSMFIDLIEENCR------MRKTSPCTDAKTPFLLAWDSVEEIV
KH.L125.4       KELQVYLDPQHTGFINYHSFLQLFESREHVTAHKWLFSTHKFNENQSPAILAWNTVEEIL
                ***   *** .**.:*   *::*:*.. :      : .*   .: ::* :***::****:

EFCAB6          HDTITRNLQAFYNMLRSYDLGDT---GRIGRNNFKKIMHVFCPFLTNAHFIKLCSKIQDI
KH.L125.4       REKLSENYRMVDKVADEYSALDTTGDGRISRTHLRRLINRYALPVSDEHFNKMWSACEGS
                ::.::.* : . ::  .*.  **   ***.*.::::::: :.  ::: ** *: *  :.

EFCAB6          GSGRILYKKLLACIGIDGPPTVSPVLVPKDQLLSEHLQKDEQQQPDLSERTKLTEDKTTL
KH.L125.4       SEGRICFARFLDNLNID--------VQPGDL---EGTSKRIQEESSEREERINEIDKADL
                ..*** : ::*  :.**        : * *    *  .*  *::..  *.     **: *

EFCAB6          --TKKMTTEEVIEKFKKCIQQQDPAFKKRFLDFSKEPNGKINVHDFKKVLEDTGMPMDDD
KH.L125.4       RHTNLLSASEVVRRLKDRMLQHDAGIRKSFLRFSRSGKDRVTKKDLRKMLDDIGLRMEDS
                  *: :::.**:.::*. : *:*..::* ** **:. :.::. :*::*:*:* *: *:*.

EFCAB6          QYALLTTKIGFEKEGMSYLDFAAGFEDPPMRGPETTPPQPPTPSKSYV----NSHFITAE
KH.L125.4       QFKELMKMLHVNGRGLLYSDFVESFQDPRTDGMGKVRRQELERAGNHIVNPTNVRYMTAQ
                *:  * . : .: .*: * **. .*:**   *  ..  *    : .::    * :::**:

EFCAB6          ECLKLFPRRLKESFRDPYSAFFKTDADRDGIINMHDLHRLLLHLLLNLKDDEFERFLGLL
KH.L125.4       ECYDQLLERMRQNFGSVRSAFYKVDDDHDGNLTMTEFRRLFDSFMFIITDQTFHELLRML
                ** . : .*:::.* .  ***:*.* *:** :.* :::**:  ::: :.*: *..:* :*

EFCAB6          GLRLSVTLNFREFQNLCEKRPWRTDEAPQRLIRPKQKVADSELACEQAHQYLVTKAKNRW
KH.L125.4       GLTKRSSLSYHDFLNKEEGHPWLKSE--HRYNRPR---SATELAADQVHQCLCLKAEQAW
                **    :*.:::* *  * :** ..*  :*  **:   : :***.:*.** *  **:: *

EFCAB6          SDLSKNFLETDNEGNGILRRRDIKNALYGFDIPLTPREFEKLWARYDTEGKGHITYQEFL
KH.L125.4       SDLAKAFQNFDADGNGIIKKKELRSVLFRFILPISHAEFNKLWSRYDEEGKGFISHQDFT
                ***:* * : * :****:::::::..*: * :*::  **:***:*** ****.*::*:*

EFCAB6          QKLGINYSPAVHRPCAEDYFNFMGHFTKPQQLQEEMKELQQSTEKAVAARDKLMDRHQDI
KH.L125.4       RKLGVGFTTGDNLVLKEMSKNKMGFF----QPQKMFCQIKKKLNIPSSHRDRFREYYSDF
                :***:.::.. :    *   * **.*    * *: : ::::. : . : **:: : :.*:

EFCAB6          SKAFTKTDQSKTNYISICKMQEVLEECGCSLTEGELTHLLNSWGVSRHDNAINYLDFLRA
KH.L125.4       DKAFRQLDKNRDGYITIADLQRVLLQLNYFLDEKQFLDLLRRLGLPTTKSKLSYFDFLRS
                .*** : *:.: .**:*..:*.** : .  * * :: .**.  *:.  .. :.*:****:

EFCAB6          VENSKSTGAQPKE---KEESMPINFATLNPQEAVRKIQEVVESSQLALSTAFSALDKEDT
KH.L125.4       IDDGRASKYGRRDVIGRESVTWQSFESLTVEKATIKLKEQVTVNYDSLNAAFRAFDRLQT
                :::.:::    ::   :*.    .* :*. ::*. *::* *  .  :*.:** *:*: :*

EFCAB6          GFVKATEFGQVLKDFCYKLTDNQYHYFLRKLRI-----HLTPYINWKYFLQNFSCFLEET
KH.L125.4       GLVKVVDFRRLLDNFCFKLTDKQFRGVLLKCRITGGSVSSNKMINWIVFLQDFSQIKDVK
                *:**..:* ::*.:**:****:*:: .* * **       .  ***  ***:** : : .

EFCAB6          ADEWAEKMPKGPPPTSPKATADRDILARLHKAVTSHYHAITQEFENFDTMKTNTISREEF
KH.L125.4       LKEWGDYVGKIAPPQSPHELPLVEVEERISEVVTARHFQISRDFADVDYAKIFVVSREDF
                 .**.: : * .** **:  .  ::  *: :.**:::. *:::* :.*  *  .:***:*

EFCAB6          RAICNRRVQILTDEQFDRLWNEMPVNAKGRLKYPDFLSRFSSETAATPMATGDSAVAQRG
KH.L125.4       RDILNRHVMRLTDDQFNRLWAKQAVNEFNNIEYREFLKRYQLHK--------DDKVKQEG
                * * **:*  ***:**:*** : .**  ..::* :**.*:. ..        *. * *.*

EFCAB6          SSVPDVSEGTRSALSLPTQELRPGSKSQSHPCTP-----ASTTVIPGTP----PLQNCDP
KH.L125.4       QTTQK-----------PCDEIQPISNQISVPRPPSRLGTSDSHRLRGNPRPVTPLVNADS
                .:. .           * :*::* *:. * * .*     :.:  : *.*    ** *.*.

EFCAB6          IESRLRKRIQGCWRQLLKECKEKDVARQGDINASDFLALVEKFNLDISKEECQQLIIKYD
KH.L125.4       AEMKVKDLVYKSWQDIQRECKKLDLEGTGTVLPDEFVGILDSFGVMLPLEDARQLMLKYD
                 * :::. :  .*::: :***: *:   * : ..:*:.:::.*.: :. *:.:**::***

EFCAB6          L-KSNGKFAYCDFIQSCVLLLKAKESSLMHRMKIQNAHKMKEAGAETPSFYSALLRIQPK
KH.L125.4       LHEQQGRFSYREFLRHFILTLKPQDEGLLKRRKIHAA-----------------------
                * :.:*:*:* :*::  :* **.::..*::* **: *

EFCAB6          IVHCWRPMRRTFKSYDEAGTGLLSVADFRTVLRQYSINLSEEEFFHILEYYDKTLSSKIS
KH.L125.4       ------------------------------------------------------------


EFCAB6          YNDFLRAFLQ
KH.L125.4       ----------


CLUSTAL format alignment by MAFFT (v7.408)


KH.L125.4       MSAVQVIERPHPGANGGLRPSSSGVVFSPRPLLSTGRNVRSLRNSTTSLARSFHVPSTSS
KH.C1.1218      MSQT--------------------------------------------------------
                ** .

KH.L125.4       GSTTVMLAANPKLTSIEVEAIIKQKLWENMEPLKQAFQLYDADRCTNVTKGEFRRVLENY
KH.C1.1218      ------------------------------------------------------------


KH.L125.4       CLPMTAEQFNALVVKIGTNPNGTISYLRFIDKFVTNSLTPNIGVARILSRPSSSNKQRKS
KH.C1.1218      ----------------------------------------------ILSRPA--------
                                                              *****:

KH.L125.4       QKFATVAKVPDEQGIDTIERFLQQKIAANLKNVIRSFKLFDYNMDGLVQRHELRRVIENY
KH.C1.1218      ------------------------------------------------------------


KH.L125.4       CLKLNDSQFNKLWSRYDFQHTGAINYRDFLRRLGVNAATAERHMRLEREKSVTDLILDRN
KH.C1.1218      ------------------------------------------------------------


KH.L125.4       KDKNKRKSMRSEMMKMEDVEKEFRKKMKNNYQNIKRALMTFDISSDGFISIDDLKAVLDN
KH.C1.1218      ------------------------------------------------------------


KH.L125.4       FVLPTSDEIFHQLMYKFEVRGTGKVSWEQFLSKFQDPQSNGNGQTIPIGQNHKVGCALDN
KH.C1.1218      ------------------------------------------------------------


KH.L125.4       SSTPDVIKKLKDHILNNFATLKQAFLAFDENRKGVIGRKDLRRIIETFAMKISDENFKEL
KH.C1.1218      ------------------------------------------------------------


KH.L125.4       QVYLDPQHTGFINYHSFLQLFESREHVTAHKWLFSTHKFNENQSPAILAWNTVEEILREK
KH.C1.1218      ------------------------------------------------------------


KH.L125.4       LSENYRMVADEYSALDTTGDGRISRTHLRRLINRYALPVSDEHFNKMWSACEGSSEGRIC
KH.C1.1218      ------------------------------------------------------------


KH.L125.4       FARFLDNLNIDVQPGDLEGTSKRIQEESSEREERRVILLILISSRSRINEIDKADLRHTN
KH.C1.1218      ------------------------------------------SQASRVGMLPRIEHPLSR
                                                          *. **:. : : :   :.

KH.L125.4       LLSASEVVRRLKDRMLQHDAGIRKSFLRFSRSGKDRVTKKDLRKMLDDIGLRMEDSQFKE
KH.C1.1218      LENPGNMSVRGVTRGSSRAGDVPKPHRLSRDAMSPRRKDLDPKMPIDSIPEGIEVVHSKD
                * ...::  *   *  .: ..: *..     : . * .. * :  :*.*   :*  : *:

KH.L125.4       LMK-MLHVNGRGLLYSDFVESFQDPRTDGMGKVRRQFRNHIVNPTNVRYMTAQECYDQLL
KH.C1.1218      PNKPTLPVFG-----NRAVMSRAESRGSNVSKM-----TSLSRPGTQTRIEIDELEMLLR
                  *  * * *     .  * *  :.* ..:.*:     . : .* .   :  :*    *

KH.L125.4       ERMRQ-NFGSVRSAFYKVDDDHDGNLTMTEFRRLFDSFM-FIITDQTFHELLRMLGLTKR
KH.C1.1218      DKIKTGGFFSVRQAFKNNDPEGKGNVTKEALMHILTSLLGRVLSSKQFHLLLKRLHLDDR
                ::::  .* ***.** : * : .**:*   : ::: *::  :::.: ** **: * * .*

KH.L125.4       SSLSYHDFLNKFEVVDKEEGHPWLNISPPYRYNRPRSATELAADQVHQCLCLKAEQAWSD
KH.C1.1218      TVVKFEEFYAQFRESVSSEYPRWLD--PVARTQTERAS--MTASQVHAQLKERAKQRFLD
                : :.:.:*  :*.   ..*   **:  *  * :  *::  ::*.***  *  :*:* : *

KH.L125.4       LAKAFQNFDADGNGIIKKKELRSVLFRFILPISHAEFNKLWSRYDEEGKGFISHQDFTRK
KH.C1.1218      LADLVPQMNPGGSERIMAPEFRNVLNRLGFFMSDAEFDKIWKKYDTSNLGVVKASSLLKK
                **. . :::..*.  *   *:*.** *: : :*.***:*:*.:** .. *.:. ..: :*

KH.L125.4       LGVGF------------TTGDNLGMLEMSKNKMGFFQPQKMFCQIKKKCEIGPQRMIPVS
KH.C1.1218      LGIEFRESKSPSASVARSTGSNKKAQVVTYDSAPPTGRSMRPSEAERQTSIDIEKWL--K
                **: *            :**.*     :: :.      .   .: ::: .*. :: :  .

KH.L125.4       YRFREYYSDFDKAFRQLDKNRDGYITIADLQRVLLQLNYFLDEKQFLDL-LRRLGLPTTK
KH.C1.1218      DKFREGFKSMRKEFKKGDLKNTGKVPRDVFRRVISQFDLFLREDSSLNMFLARCGLP-EH
                 :*** :..: * *:: * :. * :.   ::**: *:: ** *.. *:: * * ***  :

KH.L125.4       SKLSYFDFLRSIDDGRASKYGR--------------RDVIGRESVTWQSFESLTVEKA--
KH.C1.1218      GEISYVDFLNKFQDRSATGMAHNILSNPKHRFNKEPRPISPKSTVTAVESKMMTLFQSDF
                .::**.***..::*  *:  .:              * :  :.:**  . : :*: ::

KH.L125.4       ---------------------------------------------TIKLKEQVTVNYDSL
KH.C1.1218      LALLGMFHKIDKHQEDVISQQEFRAAIESRFQLEMNDDEFSHFLEQVPLNEDGAVKYPEF
                                                              : *:*: :*:* .:

KH.L125.4       NAAF-------------------------------RAFDRLQTGLVKVV---------DF
KH.C1.1218      MAQFDTKQGAKSLWDGKSVVSKAAVFVPTSDKPKERTVDDLHSILRVIVRNDMAKLEGEF
                 * *                               *:.* *:: *  :*         :*

KH.L125.4       RRLLDNFCFKLTDKQFRGVLLKCRITGGSVSSNKMINWIVFLQD----FSQIKDVK----
KH.C1.1218      RQLDEYNSGKLTQEMMFQLLSKMHIVPSVTRGEIRRLWETFIVNKNRTFSFLQFVRHYGY
                *:* :  . ***:: :  :* * :*. . . .:    * .*: :    ** :: *:

KH.L125.4       -LKEWGDYVGKIAPPQSPHELPLVE----------VEERISEVVTARHFQISRDFADVDY
KH.C1.1218      SLKSAAFPNAKIAPPQRGDNDFMIRSRKLNCAADMLEDSLRAKVDYLWEDLRREFVEMDP
                 **. .   .******  .:  ::.          :*: :   *     :: *:*.::*

KH.L125.4       AKIFVVSREDFRDILNRHVMRLTDDQXR-LWAKQAVNEFNNIEYREFLKRYQLHKDDKVK
KH.C1.1218      YHTGFVSRDEFRDVLMELCVHLTNHEAEIICNKFETNTDGRVSYVEFLRPFAQRR--QLW
                 :  .***::***:* .  ::**:.: . :  *  .*  ..:.* ***: :  ::  ::

KH.L125.4       QEGQTTQKPCDEIQPISNQISVPRPPSRLVSIMGNPRPVTPLVNADSAEMKVKDLVYKSW
KH.C1.1218      KEGNNMHS------------ILTHPQAELPGSITAGKPTKGL---EAVTSKLKQQLAGDW
                :**:. :.             :.:* :.* . :   :*.. *   ::.  *:*: :  .*

KH.L125.4       QDIQRECKKLDLEGTGTVLPDEFVGILDSFGVMLPLEDARQLMLKYDLHEQQGRFSYREF
KH.C1.1218      RTLRRAFKKMDVSADGMLTLPEFRSVLRLCNVVLDEDEVYHVLTQYD-KDLSGKLDYKKF
                : ::*  **:*:.. * :   ** .:*   .*:*  ::. ::: :** :: .*::.*::*

KH.L125.4       LRHFILTLKPQDEGLLKRRKIHAAKLPVDTGFCKLELFFVSFRSIYQRKISTTWCLSTTT
KH.C1.1218      L----------TENL----------------------------SRPSSKLSGV-------
                *           *.*                            *  . *:* .

KH.L125.4       R
KH.C1.1218      -
CLUSTAL format alignment by MAFFT (v7.408)


EFCAB6          MCKMAIIPDWLRSHPHTRKFTHSRPHS-----SPCRVYSRNGSPNKFRSSSTT-------
KH.L125.4       MSAVQVI-----ERPHPGANGGLRPSSSGVVFSPRPLLSTGRNVRSLRNSTTSLARSFHV
KH.C1.1218      MSQTIL------SRP---------------------------------------------
                *.   :      .:*

EFCAB6          -----------AVANPTLSSLDVKRILFQKITDRGDELQKAFQLLDTGQNLTVSKSELRR
KH.L125.4       PSTSSGSTTVMLAANPKLTSIEVEAIIKQKLWENMEPLKQAFQLYDADRCTNVTKGEFRR
KH.C1.1218      ------------------------------------------------------------


EFCAB6          IITDFLMPLTREQFQDVLAQIPLSTSGTVPYLAFLSRFGGIDLYIN-GIKRGGGNEMNCC
KH.L125.4       VLENYCLPMTAEQFNALVVKIGTNPNGTISYLRFIDKFVTNSLTPNIGVARILSRPSSSN
KH.C1.1218      ------------------------------------------------------------


EFCAB6          RTLRE-----------------LEIQVGEKVFKNIKTVMKAFELIDVNKTGLVRPQELRR
KH.L125.4       KQRKSQKFATVAKVPDEQGIDTIERFLQQKIAANLKNVIRSFKLFDYNMDGLVQRHELRR
KH.C1.1218      ------------------------------------------------------------


EFCAB6          VLETFCMKLRDEEYEKFSKHYNIHKDTAVDYNVFLKNLSINNDLNLRYCMGNQEVSLENQ
KH.L125.4       VIENYCLKLNDSQFNKLWSRYDFQHTGAINYRDFLRRLGVNAATAERHMRLEREKSVTDL
KH.C1.1218      ------------------------------------------------------------


EFCAB6          QAKNSKKERLLGSASSEDIWRNYSLDEIERNFCLQLSKSYEKVEKALSAGDPCKGGYVSF
KH.L125.4       ILDRNKDKNKRKSMRSEMM----KMEDVEKEFRKKMKNNYQNIKRALMTFDISSDGFISI
KH.C1.1218      ------------------------------------------------------------


EFCAB6          NYLKIVLDTFVYQIPRRIFIQLMKRFGLKATTKINWKQFLTSFHEPQGLQVSSKGPLTKR
KH.L125.4       DDLKAVLDNFVLPTSDEIFHQLMYKFEVRGTGKVSWEQFLSKFQDPQSNGNGQTIPIGQN
KH.C1.1218      ------------------------------------------------------------


EFCAB6          NSIN-SRNESHKENIITKLFRHTEDHSASLKKALLIINTKPDGPITREEFRYILNCMAVK
KH.L125.4       HKVGCALDNSSTPDVIKKLKDHILNNFATLKQAFLAFDENRKGVIGRKDLRRIIETFAMK
KH.C1.1218      ------------------------------------------------------------


EFCAB6          LSDSEFKELMQMLDPGDTGVVNTSMFIDLIEENCR------MRKTSPCTDAKTPFLLAWD
KH.L125.4       ISDENFKELQVYLDPQHTGFINYHSFLQLFESREHVTAHKWLFSTHKFNENQSPAILAWN
KH.C1.1218      ------------------------------------------------------------


EFCAB6          SVEEIVHDTITRNLQAFYNMLRSYDLGDTGRIGRNNFKKIMHVFCPFLTNAHFIKLCSKI
KH.L125.4       TVEEILREKLSENYRMVADEYSALDTTGDGRISRTHLRRLINRYALPVSDEHFNKMWSAC
KH.C1.1218      ------------------------------------------------------------


EFCAB6          QDIGSGRILYKKLLACIGIDGPPTVSPVLVPKDQLLSEHLQKDEQQQPD--------LSE
KH.L125.4       EGSSEGRICFARFLDNLNIDVQPG-------DLEGTSKRIQEESSEREERRVILLILISS
KH.C1.1218      ---------------------------------------------------------ASQ
                                                                          *.

EFCAB6          RTKLTE-DKTTL--TKKMTTEEVIEKFKKCIQQQDPAFKKRFLDFSKEPNGKINVHDFKK
KH.L125.4       RSRINEIDKADLRHTNLLSASEVVRRLKDRMLQHDAGIRKSFLRFSRSGKDRVTKKDLRK
KH.C1.1218      ASRVGMLPRIEHPLSRLENPGNMSVRGVTRG-------SSRAGDVPKPHRLSRDAMSPRR
                 :::    :     :.  .. ::  :             .    ..:  .      . ::

EFCAB6          VLEDTGMPMDDDQYA-LLTTKIGFEKEGMSYLDFAAGFEDPPMRGPETTPPQPPTPSKSY
KH.L125.4       MLDDIGLRMEDSQFK-ELMKMLHVNGRGLLYSDFVESFQDPRTDGMGKVRRQFRNHIVNP
KH.C1.1218      KDLDPKMPIDSIPEGIEVVHSKDPNKPTLPVFGNRAVMSRAESRGSNVSK---MTSLSRP
                   *  : ::.      :      :   :   .    :. .   *         .

EFCAB6          VNSHFITAEECLKLFPRRLKE-SFRDPYSAFFKTDADRDGIINMHDLHRLLLHLL-LNLK
KH.L125.4       TNVRYMTAQECYDQLLERMRQ-NFGSVRSAFYKVDDDHDGNLTMTEFRRLFDSFM-FIIT
KH.C1.1218      GTQTRIEIDELEMLLRDKIKTGGFFSVRQAFKNNDPEGKGNVTKEALMHILTSLLGRVLS
                 .   :  :*    :  :::  .* .  .** : * : .* :.   : :::  ::   :.

EFCAB6          DDEFERFLGLLGLRLSVTLNFREFQNLC------EKRPWRTDEAPQRLIRPKQKVADSEL
KH.L125.4       DQTFHELLRMLGLTKRSSLSYHDFLNKFEVVDKEEGHPWLNISPPYRYNRPR---SATEL
KH.C1.1218      SKQFHLLLKRLHLDDRTVVKFEEFYAQFRESVSSEYPRWLD--PVARTQTER-----ASM
                .. *. :*  * *     :.:.:*          *   *    .  *    :     :.:

EFCAB6          ACEQAHQYLVTKAKNRWSDLSKNFLETDNEGNGILRRRDIKNALYGFDIPLTPREFEKLW
KH.L125.4       AADQVHQCLCLKAEQAWSDLAKAFQNFDADGNGIIKKKELRSVLFRFILPISHAEFNKLW
KH.C1.1218      TASQVHAQLKERAKQRFLDLADLVPQMNPGGSERIMAPEFRNVLNRLGFFMSDAEFDKIW
                :..*.*  *  :*:: : **:. . : :  *.  :   :::..*  : : ::  **:*:*

EFCAB6          ARYDTEGKGHITYQEFLQKLGINY------------SPAVHRPCAEDYFNFMGHFTKPQQ
KH.L125.4       SRYDEEGKGFISHQDFTRKLGVGF------------TTGDNLGMLEMSKNKMGFFQPQKM
KH.C1.1218      KKYDTSNLGVVKASSLLKKLGIEFRESKSPSASVARSTGSNKKAQVVTYDSAPPTGRSMR
                 :** .. * :. ..: :***: :            :.. :        :

EFCAB6          LQEEMKELQQSTEKAVAARDKLMDRHQDISKAFTKTDQSKTNYISICKMQEVLEECGCSL
KH.L125.4       FCQIKKKCEIGPQRMIPVSYRFREYYSDFDKAFRQLDKNRDGYITIADLQRVLLQLNYFL
KH.C1.1218      PSEAERQTSIDIEKWL--KDKFREGFKSMRKEFKKGDLKNTGKVPRDVFRRVISQFDLFL
                  :  :: . . :: :    :: : ...: * * : * .. . :.   ::.*: : .  *

EFCAB6          TEGELTHL-LNSWGVSRHDNAINYLDFLRAVENSKSTG------AQPKE---KEESMPIN
KH.L125.4       DEKQFLDL-LRRLGLPTTKSKLSYFDFLRSIDDGRASK------YGRRDVIGRESVTWQS
KH.C1.1218      REDSSLNMFLARCGLPEH-GEISYVDFLNKFQDRSATGMAHNILSNPKH---RFNKEPRP
                 * .  .: *   *:.   . :.*.***. .::  ::          :.   : .

EFCAB6          FATLNPQEAVR-KIQEVVESSQLALSTAFSALDKEDTGFVKATEFGQVLKD-FCYKLTDN
KH.L125.4       FESLTVEKATI-KLKEQVTVNYDSLNAAFRAFDRLQTGLVKVVDFRRLLDN-FCFKLTDK
KH.C1.1218      ISPKSTVTAVESKMMTLFQSDFLALLGMFHKIDKHQEDVISQQEFRAAIESRFQLEMNDD
                : . .   *.  *:   .  .  :*   *  :*: : ..:.  :*   :.. *  ::.*.

EFCAB6          QYHYFLRKLRI-----HLTPYINWKYFLQNFSCFLEETADEWAEKMPKG-----------
KH.L125.4       QFRGVLLKCRITGGSVSSNKMINWIVFLQDFSQIKDVKLKEWGDYVGKI-----------
KH.C1.1218      EFSHFLEQVPL-----NEDGAVKYPEFMAQFD------TKQGAKSLWDGKSVVSKAAVFV
                ::  .* :  :          :::  *: :*.       .: .. : .

EFCAB6          PPPTSPKATADRDILARLHKAVTSHYHAITQEFENFDTMKTNTISREEFRAICNRR--VQ
KH.L125.4       APPQSPHELPLVEVEERISEVVTARHFQISRDFADVDYAKIFVVSREDFRDILNRH--VM
KH.C1.1218      PTSDKPKERTVDDLHSILRVIVRNDMAKLEGEFRQLDEYNSGKLTQEMMFQLLSKMHIVP
                ... .*:  .  ::   :   *      :  :* :.*  :   :::* :  : .:   *

EFCAB6          ILTDEQFDRLWNEMPVNAKGRLKYPDFLSRFSSETAATPMATGDSAVAQRGSSVPDVSEG
KH.L125.4       RLTDDQ-XRLWAKQAVNEFNNIEYREFLKRYQLHK--------DDKVKQEGQTTQK----
KH.C1.1218      SVTRGEIRRLWETFIVNKNRTFSFLQFVRHYGYSLKSAAFPNAKIAPPQRGDNDFMI---
                 :*  :  ***    **    :.: :*: ::            .    *.*..

EFCAB6          TRSALSLPTQELRPGSKSQSHPCTPASTTVIPGTP----PLQNCDPIESRLRKRIQGCWR
KH.L125.4       -------PCDEIQPISNQISVPRPPSRLVSIMGNPRPVTPLVNADSAEMKVKDLVYKSWQ
KH.C1.1218      ----------------RSRKLNCA-------------------ADMLEDSLRAKVDYLWE
                                .. .   .                   .*  *  ::  :   *.

EFCAB6          QLLKECKEKDVARQGDINASDFLALVEKFNLDISKEECQQLIIKYDL-KSNGKFAYCDFI
KH.L125.4       DIQRECKKLDLEGTGTVLPDEFVGILDSFGVMLPLEDARQLMLKYDLHEQQGRFSYREFL
KH.C1.1218      DLRREFVEMDPYHTGFVSRDEFRDVLMELCVHLTNHEAEIICNKFET-NTDGRVSYVEFL
                :: :*  : *    * :  .:*  :: .: : :. .:.. :  *::  : :*:.:* :*:

EFCAB6          QSCVLLLKA-KESSLMHRMKIQNAHKM---KEAGAETPSFYSALLR--IQPKIVHCWRPM
KH.L125.4       RHFILTLKP-QDEGLLKRRKIHAAKLP---VDTGFCKLELFFVSFRSIYQRKISTTW---
KH.C1.1218      RPFAQRRQLWKEGNNMHSILTHPQAELPGSITAGKPTKGLEAVTSK--LKQQLAGDWRTL
                :      :  :: . ::    :          :*  .  :  .  :   : ::   *

EFCAB6          RRTFKSYDEAGTGLLSVADFRTVLRQYSINLSEEEFFHILEYYDKTLSSKISYNDFLRAF
KH.L125.4       ---------------------------------------------CLSTTTR--------
KH.C1.1218      RRAFKKMDVSADGMLTLPEFRSVLRLCNVVLDEDEVYHVLTQYDKDLSGKLDYKKFLTEN
                                                              ** .

EFCAB6          LQ---------
KH.L125.4       -----------
KH.C1.1218      LSRPSSKLSGV
```

***Human Fibronectin1 (FN1) aligned to Ciona Fibronectin-related (Fn-rel)***

```
Heparin I binding region (FNI repeats 1-5)
Fibrillin/collagen-binding region

FN1-Hs      MLRGPGPGLLLLAVQCLGTAVPSTGASKSKRQAQQMVQPQSPVAVSQSKPGCYDNGKHYQ	60
Fn-rel      -----------------------------MRRLLLVFLFLVTIFASEVKGRCRFRGTYYR	31
                                          *:   :.     : .*: *  *  .*.:*:

FN1-Hs      INQQWERTYLGNALVCTCYGGSRGFNCE--SKPEAEETCFDKYTGNTYRVGDTYERPKDS	118
Fn-rel      SNQRWDFQVGNKNFECRCTRTGSP-HCEWSVVRVVVRRCLDS-TRRSRELGERWEVQRNN	89
             **:*:    .: : * *   .   :**      . . *:*. * .: .:*: :*  ::.

FN1-Hs      MIWDCTCIGAG---RGRISCTIANRCHEGGQSYKIGDTWRRPHETGGYMLECVCLGNGKG	175
Fn-rel      RTLDCACQENPTNHHYQITCSGKNRCHQSGISRQRGDEWTYNDA-TGLVMRCQCLGRERV	148
               **:*       : :*:*:  ****:.* * : ** *   .   * ::.* ***. :

FN1-Hs      EWTCKPIAEKCFDHAAGTSYVVGETWEKPYQGWMMVDCTCLGEGSGRITCTSRNRCNDQD	235
Fn-rel      --TCHANVRKDIPSPTPEAQVV--------NNYIVYGKTVLR-----PSLATLQACDNGH	193
              **:  ..* :   :  : **        :.::: . * *       : :: : *:: .

FN1-Hs      TRTSYRIGDTWSKK----DNRGNLLQCICTGNGRGEWKCERHTSVQTTSSGSGPFTDVRA	291
Fn-rel      --RVVTSGFTWNQTRDVTDTQYTIEQCVCRNAII------------------------RC	227
                   * **.:.    *.: .: **:* .                           *.

FN1-Hs      AVYQPQPHPQPPPYGHCVTDSGVVYSVGMQWLKTQGNK---QMLCTCLGNGVS-CQETAV	347
Fn-rel      IPYI----ISRRREPHCLTEDGEVIDVGQSWIKVHQSHENIRWRCVCASHGQRDCRDIGY	283
              *      .     **:*:.* * .** .*:*.: .:   :  *.* .:*   *:: .

FN1-Hs      TQTYGGNSNGEPCVLPFTYNGRTFYSCTTEGR--QDGHLWCSTTSNYEQDQKYSFCTDHT	405
Fn-rel      CPTPTPPI-----------NGA--IVCVSAGKAGPKQTIFCKPMCL----QNYDFHNRHR	326
              *                **     *.: *:   .  ::*.  .     *:*.* . *

FN1-Hs      VLVQTRGGNSNGALCHFPFLYNNHNYTDCTSEGRRDNMKWCGTTQNYDADQKFGFCPMAA	465
Fn-rel      ------------------------YYRVWEVCSHVTLHRWSGGY--FEDALLLGKCTRPR	360
                                     *      .:    :*.*    ::    :* *

FN1-Hs      HEEICTTNEGVMYRIGDQWDKQHDMGHMMRCTCVGNGRGEWTCIAYSQLRDQCIVDDITY	525
Fn-rel      TP-LRG-GPQSLYLQT--------------DDCLSLTRD---------------------	383
               :   .   :*                   *:.  *.

FN1-Hs      NVNDTFHKRHEEGHMLNCTCFGQGRGRWKCDPVDQCQDSETGTFYQIGDSWEKYVHGVRY	585
Fn-rel      ---------------------------------------------QIHDIEQSFVQSLRL	398
                                                         ** *  :.:*:.:*

FN1-Hs      QCYCYGR-GIGEWHCQPLQTYPSSSGPVEVFITETPSQPNSHPIQWNAPQPSHISKYILR	644
Fn-rel      NDLCGTKCQVSLFKCGERSPLPE-------------------------------------	421
            :  *  :  :. ::*   .  *.

FN1-Hs      WRPKNSVGRWKEATIPGHLNSYTIKGLKPGVVYEGQLISIQQYGHQEVTRFDFTTTSTST	704
Fn-rel      ------------------------------------------------------------	421


FN1-Hs      PVTSNTVTGETTPFSPLVATSESVTEITASSFVVSWVSASDTVSGFRVEYELSEEGDEPQ	764
Fn-rel      ------------------------------------------------------------	421


FN1-Hs      YLDLPSTATSVNIPDLLPGRKYIVNVYQISEDGEQSLILSTSQTTAPDAPPDTTVDQVDD	824
Fn-rel      ------------------------------------------------------------	421


FN1-Hs      TSIVVRWSRPQAPITGYRIVYSPSVEGSSTELNLPETANSVTLSDLQPGVQYNITIYAVE	884
Fn-rel      ------------------------------------------------------------	421


FN1-Hs      ENQESTPVVIQQETTGTPRSDTVPSPRDLQFVEVTDVKVTIMWTPPESAVTGYRVDVIPV	944
Fn-rel      ------------------------------------------------------------	421


FN1-Hs      NLPGEHGQRLPISRNTFAEVTGLSPGVTYYFKVFAVSHGRESKPLTAQQTTKLDAPTNLQ	1004
Fn-rel      ------------------------------------------------------------	421


FN1-Hs      FVNETDSTVLVRWTPPRAQITGYRLTVGLTRRGQPRQYNVGPSVSKYPLRNLQPASEYTV	1064
Fn-rel      ------------------------------------------------------------	421


FN1-Hs      SLVAIKGNQESPKATGVFTTLQPGSSIPPYNTEVTETTIVITWTPAPRIGFKLGVRPSQG	1124
Fn-rel      ------------------------------------------------------------	421


FN1-Hs      GEAPREVTSDSGSIVVSGLTPGVEYVYTIQVLRDGQERDAPIVNKVVTPLSPPTNLHLEA	1184
Fn-rel      ------------------------------------------------------------	421


FN1-Hs      NPDTGVLTVSWERSTTPDITGYRITTTPTNGQQGNSLEEVVHADQSSCTFDNLSPGLEYN	1244
Fn-rel      ------------------------------------------------------------	421


FN1-Hs      VSVYTVKDDKESVPISDTIIPAVPPPTDLRFTNIGPDTMRVTWAPPPSIDLTNFLVRYSP	1304
Fn-rel      ------------------------------------------------------------	421


FN1-Hs      VKNEEDVAELSISPSDNAVVLTNLLPGTEYVVSVSSVYEQHESTPLRGRQKTGLDSPTGI	1364
Fn-rel      ------------------------------------------------------------	421


FN1-Hs      DFSDITANSFTVHWIAPRATITGYRIRHHPEHFSGRPREDRVPHSRNSITLTNLTPGTEY	1424
Fn-rel      ------------------------------------------------------------	421


FN1-Hs      VVSIVALNGREESPLLIGQQSTVSDVPRDLEVVAATPTSLLISWDAPAVTVRYYRITYGE	1484
Fn-rel      ------------------------------------------------------------	421


FN1-Hs      TGGNSPVQEFTVPGSKSTATISGLKPGVDYTITVYAVTGRGDSPASSKPISINYRTEIDK	1544
Fn-rel      ------------------------------------------------------------	421


FN1-Hs      PSQMQVTDVQDNSISVKWLPSSSPVTGYRVTTTPKNGPGPTKTKTAGPDQTEMTIEGLQP	1604
Fn-rel      ------------------------------------------------------------	421


FN1-Hs      TVEYVVSVYAQNPSGESQPLVQTAVTNIDRPKGLAFTDVDVDSIKIAWESPQGQVSRYRV	1664
Fn-rel      ------------------------------------------------------------	421


FN1-Hs      TYSSPEDGIHELFPAPDGEEDTAELQGLRPGSEYTVSVVALHDDMESQPLIGTQSTAIPA	1724
Fn-rel      ------------------------------------------------------------	421


FN1-Hs      PTDLKFTQVTPTSLSAQWTPPNVQLTGYRVRVTPKEKTGPMKEINLAPDSSSVVVSGLMV	1784
Fn-rel      ------------------------------------------------------------	421


FN1-Hs      ATKYEVSVYALKDTLTSRPAQGVVTTLENVSPPRRARVTDATETTITISWRTKTETITGF	1844
Fn-rel      ------------------------------------------------------------	421


FN1-Hs      QVDAVPANGQTPIQRTIKPDVRSYTITGLQPGTDYKIYLYTLNDNARSSPVVIDASTAID	1904
Fn-rel      ------------------------------------------------------------	421


FN1-Hs      APSNLRFLATTPNSLLVSWQPPRARITGYIIKYEKPGSPPREVVPRPRPGVTEATITGLE	1964
Fn-rel      ------------------------------------------------------------	421


FN1-Hs      PGTEYTIYVIALKNNQKSEPLIGRKKTDELPQLVTLPHPNLHGPEILDVPSTVQKTPFVT	2024
Fn-rel      ------------------------------------------------------------	421


FN1-Hs      HPGYDTGNGIQLPGTSGQQPSVGQQMIFEEHGFRRTTPPTTATPIRHRPRPYPPNVGEEI	2084
Fn-rel      ------------------------------------------------------------	421


FN1-Hs      QIGHIPREDVDYHLYPHGPGLNPNASTGQEALSQTTISWAPFQDTSEYIISCHPVGTDEE	2144
Fn-rel      ------------------------------------------------------------	421


FN1-Hs      PLQFRVPGTSTSATLTGLTRGATYNVIVEALKDQQRHKVREEVVTVGNSVNEGLNQPTDD	2204
Fn-rel      ------------------------------------------------------------	421


FN1-Hs      SCFDPYTVSHYAVGDEWERMSESGFKLLCQCLGFGSGHFRCDSSRWCHDNGVNYKIGEKW	2264
Fn-rel      ------------------------------------------------------------	421


FN1-Hs      DRQGENGQMMSCTCLGNGKGEFKCDPHEATCYDDGKTYHVGEQWQKEYLGAICSCTCFGG	2324
Fn-rel      ------------------------------------------------------------	421


FN1-Hs      QRGWRCDNCRRPGGEPSPEGTTGQSYNQYSQRYHQRTNTNVNCPIECFMPLDVQADREDS	2384
Fn-rel      ------------------------------------------------------------	421


FN1-Hs      RE	2386
Fn-rel      --	421
```
